# Modeling Epithelial Morphogenesis and Cell Rearrangement during Zebrafish Epiboly: Tissue Deformation, Cell-Cell Coupling, and the Mechanical Response to Stress

**DOI:** 10.1101/2025.02.12.637977

**Authors:** Sharon B. Minsuk, T. J. Sego, David M. Umulis, Mary C. Mullins, James A. Glazier

**Affiliations:** Dept. of Intelligent Systems Engineering and Biocomplexity Institute, Indiana University, Bloomington, IN; Dept. of Medicine, University of Florida, Gainesville, FL; Weldon School of Biomedical Engineering, Purdue University, West Lafayette, IN; Dept. of Cell and Developmental Biology, University of Pennsylvania School of Medicine, Philadelphia, PA

## Abstract

Morphogenesis in early development involves complex and extreme deformations in response to intra- and intercellular forces. Zebrafish epiboly, the spreading of the blastoderm to cover and engulf the large yolk cell, is a key early event that sets the stage for the establishment of the body plan, but the way the forces driving expansion are generated and mediated is poorly understood. The enveloping layer (EVL), the thin squamous outer epithelium of the blastoderm, plays a central role. Forces generated in the yolk cell are transmitted through tight junctions to the marginal EVL cells, and then propagate through the rest of the EVL. To understand mechanisms of force generation and transduction during epiboly, we first need a mechanical model of the EVL capable of responding to such forces and undergoing the drastic deformation of epiboly. The expanding EVL more than doubles its surface area and experiences significant shear as it deforms from a thin cap at one pole to become a complete sphere, necessarily requiring extensive internal rearrangement. We constructed an agent-based model of the EVL and its response to exogenous forces using the center-based simulation framework, Tissue Forge. Our model captures the large viscoelastic deformation of the EVL by cell rearrangement, and incorporates algorithmic strategies to accommodate these dynamic changes while maintaining tissue cohesion. Features observed in living embryos, such as the straightening of the initially ragged leading edge, also emerge in the model. We identified two key components required for realistic epiboly in the model: first, a mechanism to enable tissue remodeling by cell rearrangement without tearing the tissue, and second, a negative feedback on the forces driving EVL expansion, to regulate and synchronize the advancement of the EVL margin. We discuss the implications of these findings for the behavior of living EVL and the mechanisms that drive epiboly.

## Introduction

Early animal development is marked by long range tissue movements with complex and extreme tissue deformations. These transformations require cells to generate and respond to intra- and intercellular forces leading to coordinated shape changes and rearrangements at both the cell and tissue levels, and ultimately to the complex structure of the adult organism. In the zebrafish *Danio rerio* and other actinopterygian (ray-finned) fish, the earliest major morphogenetic movement is epiboly, during which the blastoderm – a layer of small cells covering the animal pole region of the single large yolk cell – spreads over the yolk surface toward the vegetal pole to entirely engulf the yolk. Epiboly begins just prior to, and overlaps, gastrulation, setting the stage for the establishment of the body plan and extension of the anterior-posterior axis. Despite the apparent simplicity of this movement, the mechanisms of force generation driving it are poorly understood.

By the end of the rapid cell proliferation stages that precede epiboly, the zebrafish embryo consists of a single large cell occupying about two thirds of the embryo volume vegetally (the yolk cell), and a mass of thousands of small cells occupying the remaining third of the volume animally (the blastodisc); the boundary between these two regions is a flat plane. During doming, the first phase of epiboly, the yolk bulges upwards under the blastodisc, which becomes a cup-shaped shell of uniform thickness (the blastoderm) capping the animal end of the yolk. The blastoderm consists of two layers: the enveloping layer (EVL), a simple, non-stratified, squamous epithelium, and the deep layer, composed of mesenchymal cells, several cells thick (Fig. 1a, 30%). The deep layer lies between the EVL and the yolk, except at the outer margin of the blastoderm where the EVL leading edge cells come into contact with the yolk cell, and are bound to it by tight junctions. At this stage this leading edge of the EVL has ragged edges; it will straighten markedly during the early part of epiboly, becoming a smooth circumferential boundary and remaining straight thereafter. A layer of syncytial nuclei within the yolk cell (the yolk syncytial layer, YSL) lies beneath the blastoderm, and its external margin (the e-YSL) in the yolk cortex is overlain by a dense band of filamentous actin (Fig. 1a, 50% and 75%). The EVL, deep cells, YSL, and actin band all undergo epiboly together [1–4].

**Fig. 1.**
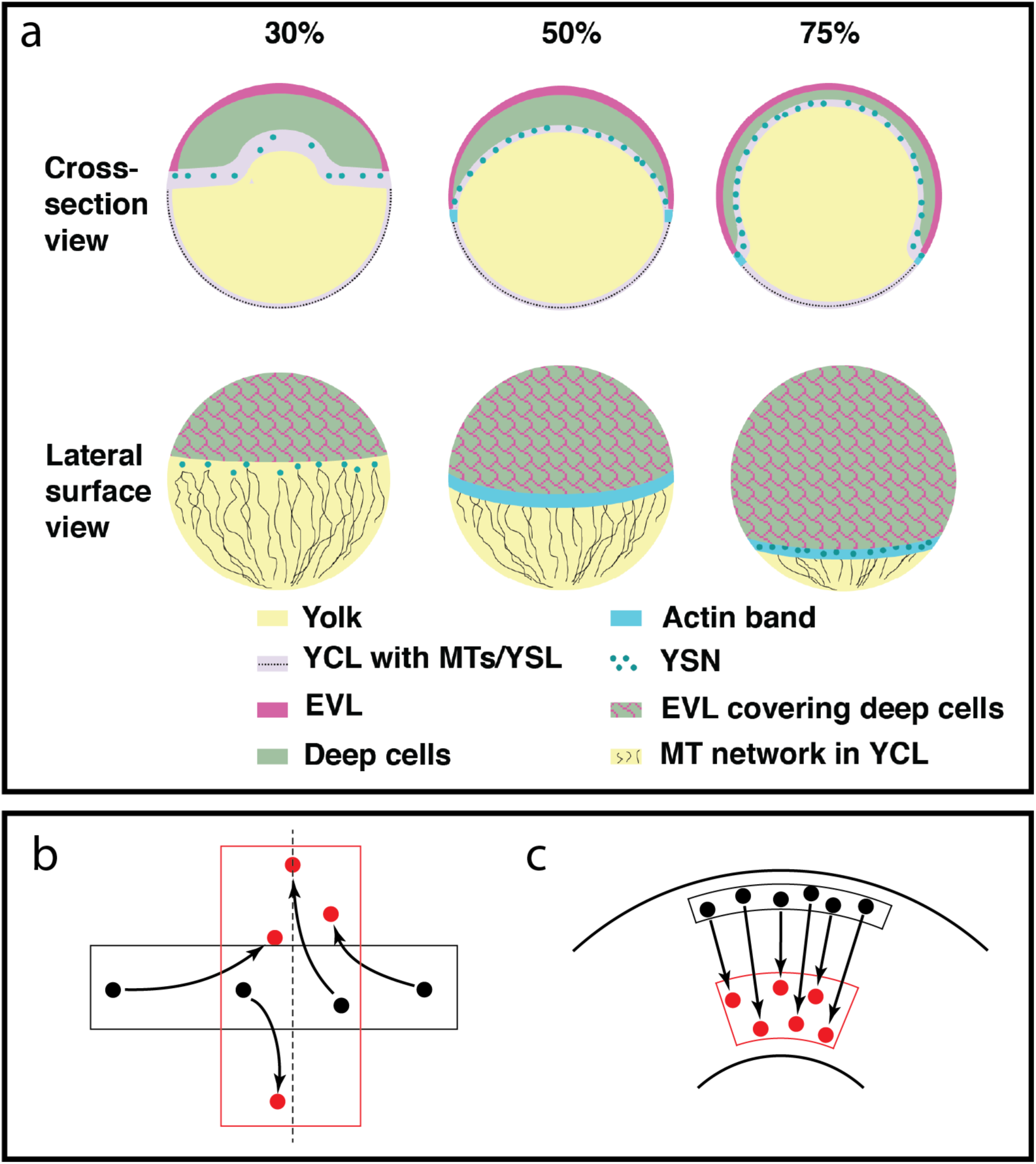
a. Schematic of epiboly progression. Epiboly begins as the outer-most epithelial sheet cells, the enveloping layer (EVL, pink outline), expand, while the yolk (yellow) domes toward the animal pole (top), and the deep cells of the blastoderm (green) radially intercalate to cover 30% of the embryo surface. At 30% epiboly, the yolk syncytial nuclei (YSN, dark green circles) reside in a yolk cytoplasmic layer (YCL) beneath the blastoderm and are associated with microtubules (black lines, surface view) that are oriented toward the vegetal pole within the very thin cortical YCL (light blue in cross-section). The EVL is a thin epithelial sheet covering the surface of the deep cells and is connected to the yolk membrane via tight junctions at the EVL leading edge. At 50% epiboly, the EVL and deep cells cover 50% of the embryo and lie more vegetally than the YSN, masking the YSN in the lateral surface view. An actin band (turquoise) forms in the YSL just vegetal to the margin at 50% epiboly, where it acts in endocytosis of the yolk cell membrane ahead of the EVL margin, and may function as a contractile mechanism in the completion of epiboly. By 75% epiboly, the YSN are largely located beneath the blastoderm, as well as vegetal to it (surface view). The exposed vegetal yolk continually decreases in circumference as epiboly progresses, until the blastoderm completely engulfs the embryo. Figure and modified legend taken from [23]. b and c: Schematic diagrams of two styles of convergence and extension movements (CE). Black-bounded regions each represent a patch of tissue at the beginning of CE, and black dots indicate the positions of several cells within that patch. Red-bounded regions and dots represent the tissue shape and cell positions after the process completes. Arrows mark the trajectories of the cells as the tissue changes shape. b: Traditional CE toward a midline (dashed line), such as during axial elongation of the zebrafish deep cell layer. c: Uniform CE around the cylindrical geometry of the zebrafish embryo, without a midline or lateral cell movement, as the circumference of the leading edge shrinks, viewed as from the vegetal pole.

Of these layers, we single out the EVL for modeling in this study because of its central role in epiboly. Spreading of the deep cells is partially dependent on adhesion to the EVL; but the dependence is not mutual, as the EVL can spread when the adhesion is disrupted [2]. Several complementary force generating mechanisms have been proposed to explain what drives EVL spreading; all of them posit an exogenous driving force that arises within the yolk cortex, acts directly on the yolk-EVL tight junctions that mechanically couple the cortex to the marginal EVL cells, and is then propagated back through the successive tiers of EVL cells, making the EVL margin a focal point of any putative force-generating mechanism. Several proposed mechanisms hinge on actomyosin-generated forces at the boundary where the leading edge of the EVL meets the yolk cell surface. EVL expansion begins when the balance of forces in the EVL and the yolk cell cortex shifts due to relaxation within the EVL, so that the yolk cortical tension dominates [5]. Thereafter, expansion continues due to ongoing generation of vegetally-oriented forces within the yolk cell [3]. The actin band in the YSL is proposed to drive endocytosis of the overlying yolk cell membrane just ahead of the EVL margin [6], which generates membrane tension driving EVL expansion [7–10]. The same band is also proposed to contribute to EVL expansion by constriction in both the animal-vegetal and the circumferential directions [9,11,12] and by an actomyosin flow-friction force-generating mechanism [12]. Contraction of actin rings at the leading edge of both the EVL and the deep cells may also contribute to expansion [6]. Finally, microtubule-dependent towing of the blastoderm may be associated with an elaborate microtubule network in the yolk cortical cytoplasm (the main driver of YSL spreading) [13]. To model the relative contributions of these putative mechanisms, we need a mechanical model of the EVL capable of responding to such forces and undergoing the drastic deformation of epiboly.

The zebrafish EVL must increase its surface area about 2.3-fold during epiboly, which it accommodates by apicobasal thinning [14]. It must also undergo considerable shear to deform from its initial spherical cap shape, to the final full sphere. The permanence of the deformation reflects the viscoelasticity of the tissue, an important feature of tissues undergoing morphogenesis in all animals. Viscoelastic tissue can deform without tearing, undergoing extensive internal reorganization reflected in either cell rearrangement or cell shape change or a combination of both. Zebrafish EVL displays both types of reorganization, but appreciable cell elongation occurs only late in epiboly, and only close to the leading edge [1,15]. Therefore the deformation during early and middle stages of epiboly is dominated by cell rearrangement. Cell rearrangement within a mechanically coupled epithelium requires the cells to slide between one another, exchanging neighbors without ever breaking the epithelial barrier [15].

Cell division, which is rapid and synchronous during cleavage stages, slows down and becomes asynchronous during epiboly [4]. In the EVL, cell division gradually slows until it becomes negligible by around 60% epiboly [14]. Epiboly can proceed, at least to some degree, even when cell division is suppressed, but division may be necessary for epiboly completion. Moreover, oriented division aligned preferentially in the animal-vegetal direction appears to play some role, by releasing tension and promoting vegetalward tissue elongation [14].

EVL stretch, deformation, cell rearrangement, and division must take place without disrupting the mechanical coupling between the EVL cells that holds the epithelial tissue together, or its physiological barrier function [15]. Some recent modeling studies have treated the spreading EVL as a uniform fluid [5,12]. That approach addresses global tissue-level properties, but cannot account for individual cell behaviors and interactions, essential for an understanding of the cellular mechanisms underlying emergent tissue-level behaviors. To that end, we used an an agent-based approach. In an agent-based model, a large collection of objects (in this case, EVL cells) is modeled by explicitly describing each individual object as an autonomous unit (“agent”), specifying the properties of each type of agent, and defining the interaction rules between them; their collective behavior arises from those properties and rules.

The transformation from spherical cap to full sphere implies certain dynamics that our model must capture. At first, as the EVL expands in surface area above the equator, the EVL expands both circumferentially (the length of the leading edge of the EVL increases as the EVL stretches over the widest part of the embryo) and longitudinally (the distance from the animal pole to the leading edge increases). But once the EVL leading edge passes the equator (50% epiboly), regions below the equator must shrink in circumference while continuing to expand vegetally, constituting convergent extension (CE) toward the vegetal pole (Fig. 1b,c). For the purposes of this study we define CE as a tissue shape change rather than the cellular mechanisms underlying it; therefore, not as a cause of morphogenesis, but as a macroscopic description of it. The term “convergent extension” in zebrafish classically refers to the narrowing of axial tissue toward the dorsal midline and its animal-vegetal elongation [16] (Fig. 1b), which takes place in the deep cells and does not involve the EVL. In contrast, the deformation of the blastoderm leading edge during epiboly (both EVL and deep cells) is a second and distinct instance of CE, representing a variation on the process that occurs as the blastoderm circumference shrinks uniformly and the tissue expands vegetally, requiring neither an identifiable midline, nor lateral migration [17] (Fig. 1c). Our model considers only epiboly and not axial development, and only the EVL, so our discussion of convergent extension refers exclusively to this second process. See Appendix for discussion of the variety of CE modalities and the justification for our terminology.

We used the Tissue Forge simulation framework [18] to build a model of zebrafish EVL epiboly, identifying critical components sufficient to reproduce the observed behaviors of living EVL listed on the right side of Table 1. We found two key components essential to the model.

**Table 1.** Model algorithm components and resulting model properties.

| Algorithm components |  |  |  |  | Simulation behaviors |  |  |  |  |
| --- | --- | --- | --- | --- | --- | --- | --- | --- | --- |
|  | Bond remodeling | Remodeling algorithm | Force regulation | Cell division | Viscoelasticity | Tissue cohesion | Synchrony | Cell rearrangement | Edge Straightness |
| **1. Living zebrafish embryos** |  |  |  |  | + | + | + | + | ↑ |
| 2. Simplistic model: fixed bonds | – |  | – | – | – | + | + | – | ↓ |
| 3. Dynamic model: bond remodeling | + | R | – | – | – | – | – | + | ↓ |
| 4. Model 1 (viscoelastic: constrained geometry) | + | EM | – | – | + | + | – | + | ↑<sup>a</sup> |
|  |  |  |  | + | + | + | – | + | ↑<sup>a</sup> |
| 5. Model 2 (regulated: force feedback) | + | EM | + | – | + | + | + | + | ↑ |
|  |  |  |  | + | + | + | + | + | ↑ |

Line 1 represents the properties of living embryos we would like our model to replicate.
Subsequent lines represent different versions of our model, including different combinations of algorithm components, and the resulting model properties.

First, a mechanism to enable controlled tissue remodeling by cell rearrangement, to allow for viscoelastic tissue deformation without tearing (Table 1, line 4). Stochastic remodeling of cell-cell mechanical coupling (cell adhesion) allows for rearrangement, and an energy-minimizing constraint on the resulting packing geometry prevents tearing. Second, a negative feedback on the forces pulling the EVL leading edge, to synchronize its advance so that all parts of it advance simultaneously (Table 1, line 5). Additionally, we observed that the initially ragged leading edge of the model EVL straightens during epiboly (Table 1, lines 4 and 5), unexpectedly reproducing, emergently and robustly, a poorly studied feature of living embryos (Table 1, line 1) [1]. In the model, the straightening occurs most quickly under conditions allowing rapid cell rearrangement and tissue fluidity, suggesting a role in living EVL for a solid-like to fluid-like transition underlying both tissue deformation and straightening of the EVL edge.

### Modeling epiboly

#### Conceptual biological model

From the biological observations described above, we identify a set of properties to build into our model, and behaviors that our model must recapitulate. Our simulation will begin at the 30% epiboly stage, so the EVL will begin as a cap covering the embryo from the animal pole down to around 8° above the equator, consisting of roughly isodiametric cells, equal in size. Neighboring cells will be mechanically coupled, as in all epithelia, so that forces acting on one cell can propagate to its neighbors and throughout the epithelium.

In response to external, vegetally directed forces acting on the leading edge cells, the EVL must expand toward the equator, increasing both in area and in the circumference of its leading edge. After passing the equator, with further tissue expansion the leading edge circumference must shrink, progressing vegetally and converging toward the vegetal pole until the EVL finally engulfs the yolk completely. All points along the leading edge must advance synchronously to the pole. In undergoing this expansion and engulfment, the EVL must undergo permanent viscoelastic deformation, without tearing, accommodated by cell rearrangement (Table 1).

Because we focus on the response of the EVL to stretch, we can treat other parts of the system as simple boundary conditions. We do not model the deep cell layer explicitly, but treat it along with the yolk cell as a single composite object representing the large embryo mass over which the EVL spreads. (We refer to it as the “yolk cell” for simplicity.) We treat the external driving force, generated from within the yolk cell, as a generic force acting on the leading edge cells, pulling them toward the vegetal pole, without specifying any particular mechanism of force generation or the detailed structure of the yolk cortical region.

Thus we conceptualize the embryo as a single roughly spherical yolk cell, covered near its animal pole by a monolayer of EVL cells. These cells can exert forces on one another, receive forces acting upon them, and respond to these forces by moving (translocating in the direction of net force) and/or stretching/compressing (changing their apical surface area, represented as a change in the distance between neighbor cell centers). We model two types of forces to represent the interactions between epithelial cells (Table 2). First, neighboring cells are held together by cell adhesion; we model this as an attractive force between a pair of neighboring, adhering cells, resisting any stretching force that would otherwise drive the centers of the two cells apart. Second, the centers of two cells (whether or not the cells are adhering) cannot approach each other arbitrarily closely, because cells resist compression. We model this resistance as a repulsive force between the cell centers. Finally, we include cell division in the model. We set out to model these component cells and forces and the dynamic interactions between them (Table 2) in such a way that the model EVL can undergo morphogenesis as described above.

**Table 2:** Model components.

| **Biological** | **Mathematical model** | **Implementation with Tissue Forge** |
| --- | --- | --- |
| ***Objects*** |  |  |
| EVL cells | Point masses arranged in a monolayer | Particles |
| Yolk cell and deep cell layer | Abstracted as a single point object | One large particle |
| ***Processes*** |  |  |
| Resistance to compression | Repulsive force between two cells under pressure, modeled as a spring | Harmonic potential: non-bonded (active whenever two cells are in proximity, as defined by cell radius) |
| Cell adhesion | Attractive force between adhering EVL cells when stretched, modeled as a spring | Harmonic potential: bonded (active when adhesion explicitly present between two cells) |
| EVL–yolk cortex mechanical coupling by tight junctions, and vegetalward pulling force | Constant, uniform force per unit length of EVL edge, applied to leading edge cells only | Vector force on discrete particles, weighted according to length of contact between EVL cell and yolk cortex |
| ***Dynamics*** |  |  |
| Exogenous (yolk-derived) force experienced by an EVL cell changes as its yolk contact length changes | Contact-length depends on relative positions of other EVL cells, and on cell rearrangement (cells can enter/leave the leading edge) | Vector force per each discrete particle dynamically recalculated each timestep |
| EVL cells modify cell-cell adhesive contacts, allowing exchange of neighbors | Stochastic activation/deactivation of attractive forces | Dynamic making and breaking of bonds |
| Cell division within plane of epithelium. | Apico-basal height of daughter cells same as parent; apical and basal surface area half that of parent. | Particle duplication with reduced cell radius equivalent to 50% reduction in surface area. |

#### Mathematical/Quantitative model

We determined the number of EVL cells, and rates and patterns of cell division, based on [14]. We optimized other physical parameters for performance by trial and error. See Methods for a detailed description, and Table 3 for a list of key configuration parameters.

**Table 3:** Key configuration parameters (constant over all simulations described, except where noted). Parameter names as specified in our python simulation code. See Methods for details.

| Parameter name | Default value | Notes |
| --- | --- | --- |
| **Initialization:** |  |  |
| epiboly_initial_num_evl_cells | 660 | Number of EVL cells at 30% epiboly [14]. Each simulation begins with a stochastically determined approximation of this value. |
| epiboly_initial_percentage | 43 | The true leading edge position at the misnamed "30% epiboly"; see Methods. |
| cell_division_cessation_percentage | 55 | Stage of epiboly progression at which cell division, if enabled, ceases [14]. |
| total_epiboly_divisions | 412 | Increase in number of EVL cells while division is active [14]. Each timestep, a stochastically generated number of cells divides, calibrated to approximate this total number of divisions by the configured cessation of cell division. |
| **Potentials and bonds:** |  |  |
| harmonic_repulsion_spring_constant | 0.5 | Repulsion between EVL cells (bound or unbound) |
| harmonic_spring_constant | 0.5 | Attraction between bound internal EVL cells |
| harmonic_edge_spring_constant | 0.5 | Attraction between bound EVL leading edge cells (constant except in the experiment in Fig. 11) |
| harmonic_yolk_evl_spring_constant | 4.0 | Attraction/repulsion between EVL and yolk cells |
| **Strength of bond angle constraint:** |  |  |
| lambda_bond_angle | 3.75 | Strength of constraint on EVL internal bond angles |
| lambda_edge_bond_angle | 4.0 | Strength of constraint on leading edge bond angles (constant except in the experiment in Fig. 10) |
| **Epiboly driving force:** |  |  |
| _force_per_unit_length | 0.7 | Applied uniformly in Model 1; weighted locally according to leading edge distance from vegetal pole in Model 2 |
| **Boolean flags to switch between versions of the model:** |  |  |
| cell_division_enabled |  | See main text |
| force_regulation_enabled |  | False for Model 1; True for Model 2 |
| bond_remodeling_enabled | True | Switched off for elastic model (no bond remodeling, Fig. 3), and during recoil test (Fig. 12) |

#### Computational model

We model cells using a center-based approach. Center-based models represent each object as a simple point mass in 3D space, which behaves as the center of mass of the 3D object it represents; modeled forces act on those centers of mass, determining the displacement of the objects. Exact sizes and shapes of the objects, or their boundaries, are not explicitly represented. We model each individual EVL cell as a single such point (Fig. 2).

**Fig. 2.**
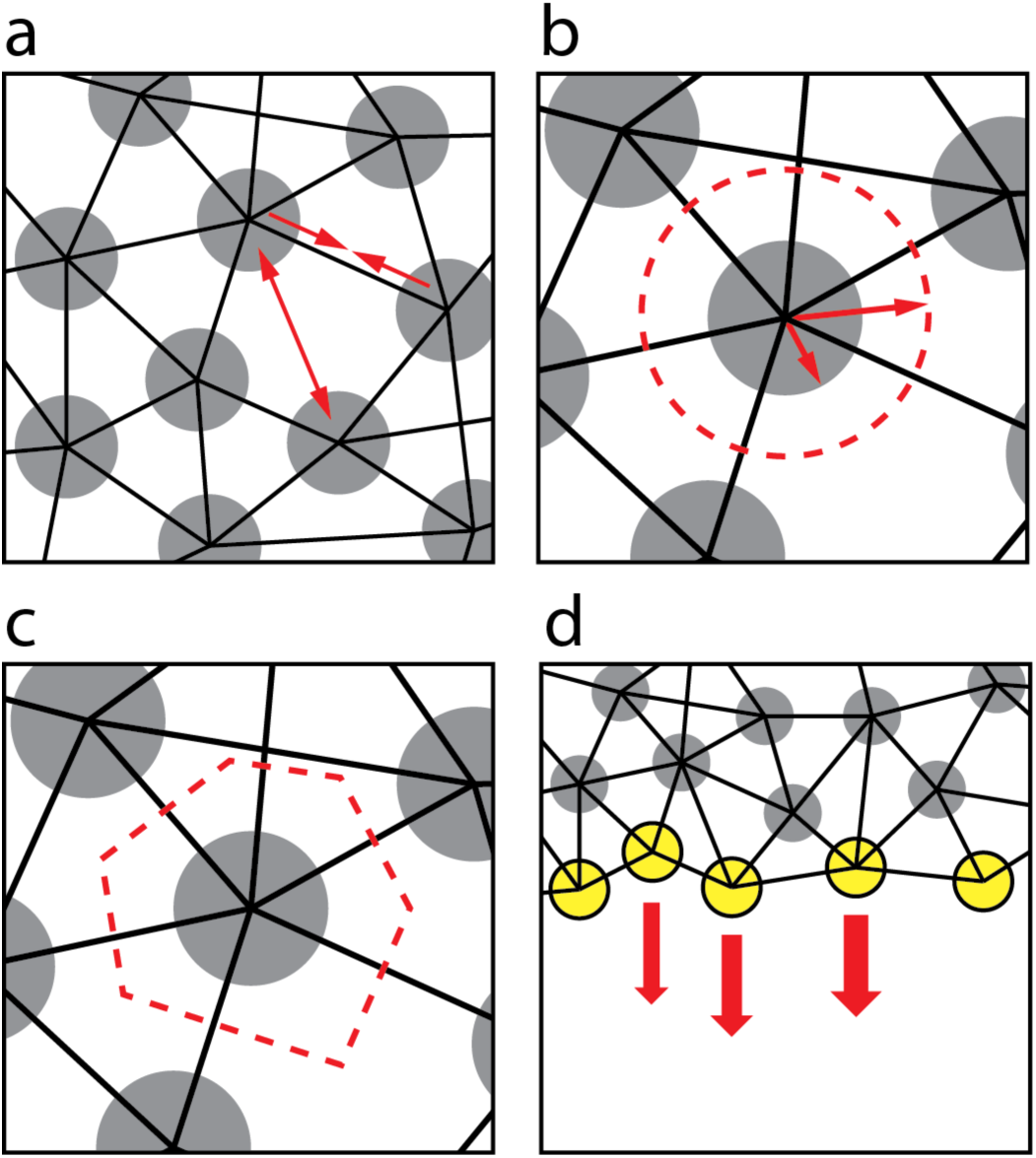
Components of the model. a. Particles (circles), bonds (black lines), and forces (red arrows). All pairs of particles, including non-bonded pairs, exert repulsive forces on one another (double-headed arrow, repulsion between two particles lacking a bond between them), representing volume exclusion. Bonded pairs, in addition to their repulsive forces, exert attractive forces on one another (pair of single-headed arrows), representing adhesion. b. Particle radius vs. cell radius. Tissue Forge particles each have a *particle* radius (short arrow), affecting only the graphical rendering and not the physics (because Tissue Forge particles represent point masses). Our model assigns to each particle, a distinct *cell* radius (long arrow). Resting lengths of each bond are set to the sum of the cell radii of the two bonded particles, thereby limiting the distance of approach between the particles. The cell radius thus represents the target or equilibrium size of the cell (dashed circle). Cell size can depart from this ideal resting size when the tissue comes under tension (i.e., if the tissue is stretched, cell surface areas over the entire EVL must on average increase). c. Cell shape and cell boundaries are not explicitly represented by the model. But cells are interpreted as epithelial, tiling the layer and leaving no gaps. Therefore the inferred cell shapes (dashed boundary) are irregular and polygonal, as typically found in epithelia. d. Exogenous forces act on the leading edge particles (yellow), pulling them vegetalward (red arrows). Arrow thickness represents magnitude of force, which for each particle is proportional to the average horizontal distance between its center and those of its two bonded leading-edge neighbors. In this way we can model a force that in living tissue is presumed uniform along the continuous EVL/yolk boundary.

We model force between two EVL cells as a spring, with a resting or equilibrium distance and a spring constant. The resting distance defines the center-to-center distance at which two cells are under neither compression nor tension, hence exerting no force on one another; the force is repulsive when the centers are closer than equilibrium, and attractive when further apart (Fig. 2a). The spring constant determines the cells’ compressibility and resistance to stretch, and the strength of cell adhesion.

We set the spring resting length for a pair of cells equal to the sum of their cell radii. We calculate an average cell radius (Fig. 2b) based on the reported number of EVL cells in live embryos [14] and the initial size of the EVL as a whole. Two cells at their equilibrium distance therefore represent cells that are just touching, and relaxed. Thus although our center-based model does not explicitly represent the location of cell boundaries, the forces between the cells reflect the cell size and determine the initial arrangement and subsequent behavior of the point masses (Fig. 2b). Deriving the spring resting length from the calculated cell radius thus endows each cell with an effective volume; and approximate cell shapes can be inferred from the locations of neighboring, adhering cells (Fig. 2c). The starting configuration is shown in Fig. 3a. To compute the external force to apply to each of the marginal EVL cells, we assume the force is generated by uniform local processes within the zebrafish yolk cell, and therefore that the force applied per unit length along the EVL margin is both spatially uniform along its circumference (Fig. 2d), and constant in magnitude over time. See Methods for details.

**Fig. 3.**
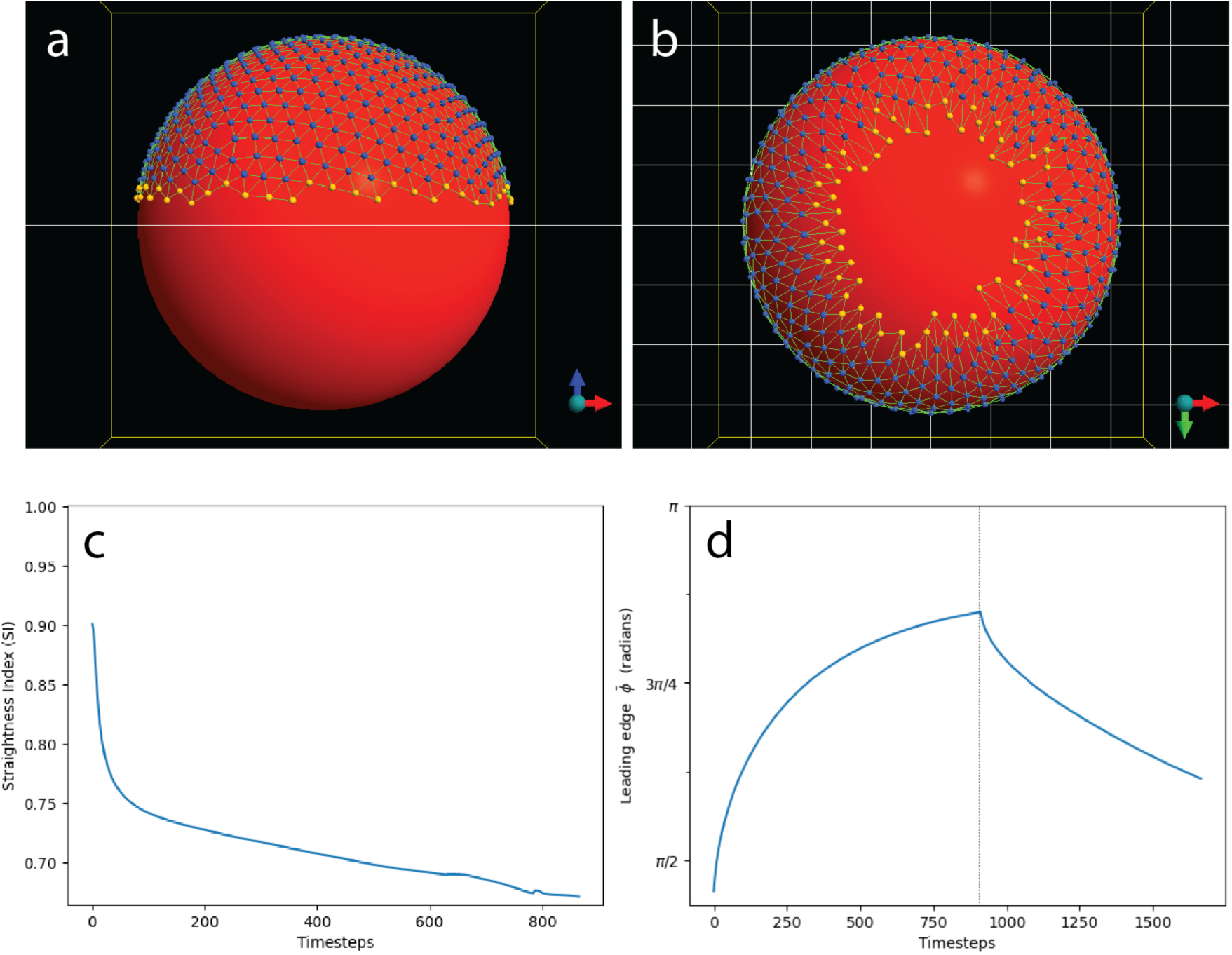
A simple preliminary model EVL (without dynamic bond remodeling). The EVL (marginal cells yellow, internal cells blue) is initialized as a cap over the top of the large yolk cell (red). Exogenous forces are applied to the marginal EVL cells only. See also Movie S1. a. The model EVL (lateral view) at initialization, representing the 30% epiboly stage. The edge of the sheet (yellow particles) is relatively straight but slightly ragged. b. The model EVL after being stretched around the yolk (vegetal view). Individual bonds, and the tissue as a whole, have undergone elastic stretch. Cell neighbor relationships are fixed (preventing any cell rearrangement). For the EVL to accommodate itself to the spherical surface of the yolk without internal reorganization, the stretched leading edge has become jagged. c. Straightness index (measuring the straightness of the leading edge; where 1 = perfectly straight and 0 = infinitely tortuous; see Methods) drastically decreases over the course of the simulation, as the tissue edge becomes highly convoluted. d. Response of the stretched cell layer to experimental disruption of forces demonstrates its elasticity. Mean polar angle of the EVL margin over time; vertical marker line indicates the end of epiboly and the start of the experiment, when the exogenously applied force is disabled. The cell layer undergoes drastic recoil.

Integration of the forces over time transforms the model components just described into a dynamic, evolving system of moving cells. How these resulting movements unfold will be further influenced by our choice of update rules: how cell adhesions change over time, how the exogenous driving force is regulated, and the presence or absence of cell division. In the Results, we describe our development and refinement of these model update rules, and our resulting working model of zebrafish epiboly.

#### Implementing the model using Tissue Forge

The Tissue Forge simulation framework [18] provides an extensive toolkit for implementing center-based models. Most important for this study are its particles (point masses), forces (defined by vectors applied to particles), and potentials (functions describing the energy relationship between two particles as a function of center-to-center distance, which determines the forces exerted by the particles on each other). Specifically, we use a harmonic potential, which behaves exactly as a spring, with a resting or equilibrium distance and a spring constant. Potentials can be applied either as bonded interactions (applied to a specified pair of particles, and forming a stable interaction until the bond is removed) or as a non-bonded interaction (global and applying to any two particles that approach each other to within a specified distance, and only while they remain in proximity). It’s worth noting that particles have a radius, which describes only the visual rendering of the particles as spheres, and does not affect the physics; it is distinct from the *cell* radius we described above (Fig. 2b). Once we define the types, instances and arrangement of these components, Tissue Forge iterates over a series of timesteps, integrating the forces and updating the particle locations at each timestep. We describe the operation of the framework and our application of it to our zebrafish model in its complete detail in the Methods.

At initialization of the model, equally sized EVL cells are stochastically positioned in a monolayer over the top of the yolk, covering an area equivalent to the 30% epiboly stage of zebrafish. Repulsive forces hold the cell centers apart, representing the incompressibility of water/cytoplasm; and attractive forces hold the cells together, representing cell adhesion and providing global mechanical coupling (Fig. 2a). As the simulation proceeds, these cell adhesions can be dynamically added and removed. An exogenous pulling force is applied uniformly along the leading edge, acting on the centers of mass of the cells, directed always tangent to the yolk surface, oriented toward the vegetal pole (Fig. 2d). (See Methods for more detail.)

## Results

### Modeling a viscoelastic, mechanically coupled cell sheet that can remain intact while deforming under tension (Model 1)

The model components so far described – if the bonded relationships between particles (the nodes and edges of a connected graph) remain fixed throughout the simulation – create an elastic cell layer, because the individual bonds are elastic (they will return to their equilibrium length when unconstrained). To demonstrate the properties of such an elastic sheet (Table 1 line 2) and contrast them with the behavior of living EVL (Table 1 line 1) and of the final model we build, Fig. 3 and Movie S1 show this initially configured structure being stretched around the sphere toward the vegetal pole, followed by the recoil of the sheet after the stretching force is removed. When the sheet of particles with this fixed topology is forced to converge toward the vegetal pole, the cells become elongated (as inferred from the elongated bonds and the sharp angles between them; Fig. 3b, Movie S1), and the leading edge inevitably becomes jagged (Fig. 3b,c) because when bonded cell neighbor relationships can not be remodeled, the increasingly crowded leading edge cells cannot be accommodated along a straight edge and the edge cannot shorten to fit the smaller embryo circumference near the pole. (The tissue would buckle into the third dimension except that the particles are held tightly to the yolk surface; see Methods.)

To capture the permanent viscoelastic deformation of the EVL and the accompanying cell rearrangement, we must provide for dynamic remodeling of the tissue geometry as it stretches (Table 2). We therefore allow for reorganization of cell-cell connectivity by applying dynamic, stochastic bond remodeling (breaking existing bonds and forming new ones). This is not intended to represent a literal biophysical description of how living EVL cells rearrange; we do not suppose that neighboring epithelial cells completely release their adhesion to one another. Our bond-breaking and re-formation is instead an abstraction of the gradual remodeling of adhesive junctional relationships required for neighbor exchange [15]. We developed a bond remodeling algorithm, described in detail in the Methods. During each simulation timestep, at each EVL cell, either a randomly selected existing bond is broken, or a new bond to a nearby cell is created (Fig. 4).

**Fig. 4.**
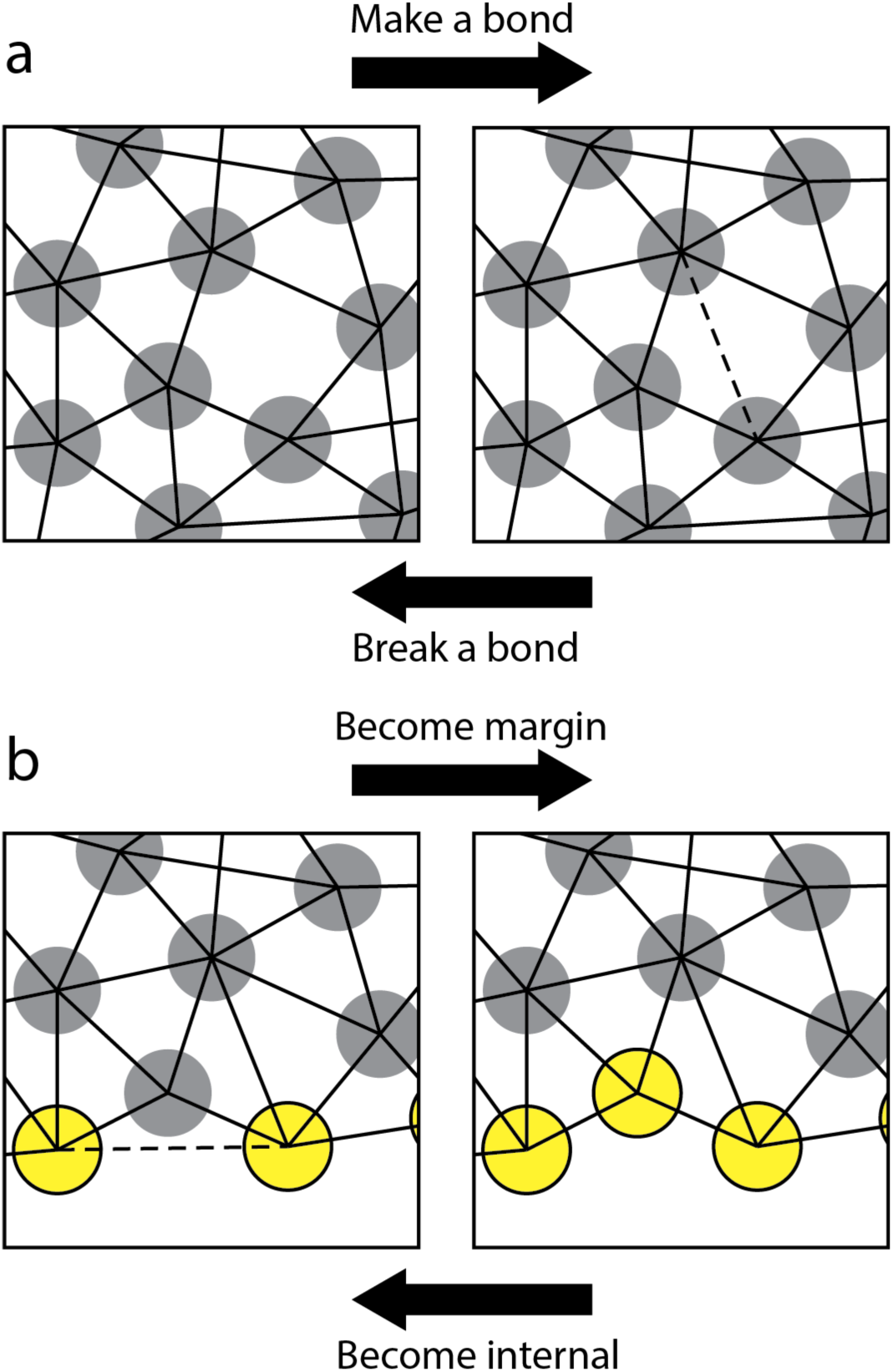
The four bond-remodeling transformations. a. Forming a new bond (proceed from left panel to right); or breaking an existing one (from right to left). b. An internal cell becomes part of the margin, requiring the breaking of a bond between its two marginal neighbors (proceed from left panel to right); or a marginal cell leaves the margin and becomes internal, as its two marginal neighbors form a bond (from right to left). Grey circles: internal EVL particles; yellow circles: EVL margin particles; solid lines: bonds connecting centers of two particles; dashed lines: bonds being formed or broken during a transformation.

We found that with this simple approach to bond remodeling, EVL integrity could not be maintained (Table 1 line 3); large tears would develop in the tissue under increasing tension. Tissue tearing occurred because bond breaking and bond formation were not inherently spatially coupled; therefore any local excess of bond breaking over bond formation could lead to net loss of connectivity. Living embryos solve the problem of allowing cells to rearrange even in a tissue under tension, releasing old neighbors and binding to new ones without the tissue disintegrating in the process [1,15], and so must we. We therefore introduced additional rules to improve tissue integrity by further limiting bond remodeling events that damage the epithelial structure. Initial attempts to impose a statistical correlation between the two types of events, to enhance their spatial coupling, were not productive. Instead, we used an energy minimization approach, defining energetically favorable transformations as those that lead to topological stability (Table 1 line 4). Specifically, for each provisional bond-breaking or bond-forming event, we applied an elastic constraint on the bond angles between pairs of adjacent neighbor bonds on each cell, defining 60° bond angles as most favorable (see Methods for details). This geometry corresponds to the natural packing of approximately equal sized, roughly isodiametric cells in a planar sheet, a consequence of cell surface tension (and treats the sheet as approximately planar since the EVL cells are small compared to the yolk cell). Bond angles along the margin prefer a roughly 180° bond angle (adjusted to take the curvature of the embryo surface into account; see Methods). Locally, the configuration is free to depart from these lowest energy states, but the greater the departure, the more energetically unfavorable the event and hence the lower the probability of the event being carried out; and if it is carried out, the greater the probability of it being quickly reversed. The algorithm is described in detail in the Methods.

Finally, we add cell division to the model. We implemented a simplified version of the cell division profile documented in living embryos by Campinho et al. [14]. In our model, cells divide with random orientation from 30% epiboly (the start of the simulation, stochastically initialized with approximately 660 cells), at regionally uniform cell division rates, and then cease dividing at 55% epiboly. We calibrate overall division rate so as to approximate a 62% increase in the number of EVL cells over the period of active division, as reported in [14]; during each simulation timestep, we determine the number of divisions stochastically. This results in a small number of cells (usually between 0 and 10) selected for division at any given timestep, at random, with the constraint that no cell is allowed to divide more than once during the simulation. Each dividing cell is split into two; we assume the apicobasal cell height of the two daughter cells after division (the local epithelial thickness) is the same as the height of the parent cell before division, and therefore the apical surface area of each daughter is set to half that of the parent. (See Methods for details.)

Together, these behavioral rules result in a robust connected EVL, able to stretch while under tension without tearing, and engulf the yolk (Fig. 5a, Movie S2; Table 1, line 4). The same result is shown with daughter cells visibly marked, in Fig. S1 and Movie S3. Cell division is not required for engulfment; epiboly progresses just as well when cell division in the model is disabled (Fig. S2, and Movie S4). In contrast with living embryos, in which all points along the EVL leading edge progress synchronously, in Model 1 all points on the leading edge do not advance in synchrony, and the leading edge does not remain circular at the latest stages; we address this with an enhanced version of the model in the next section.

**Fig. 5:**
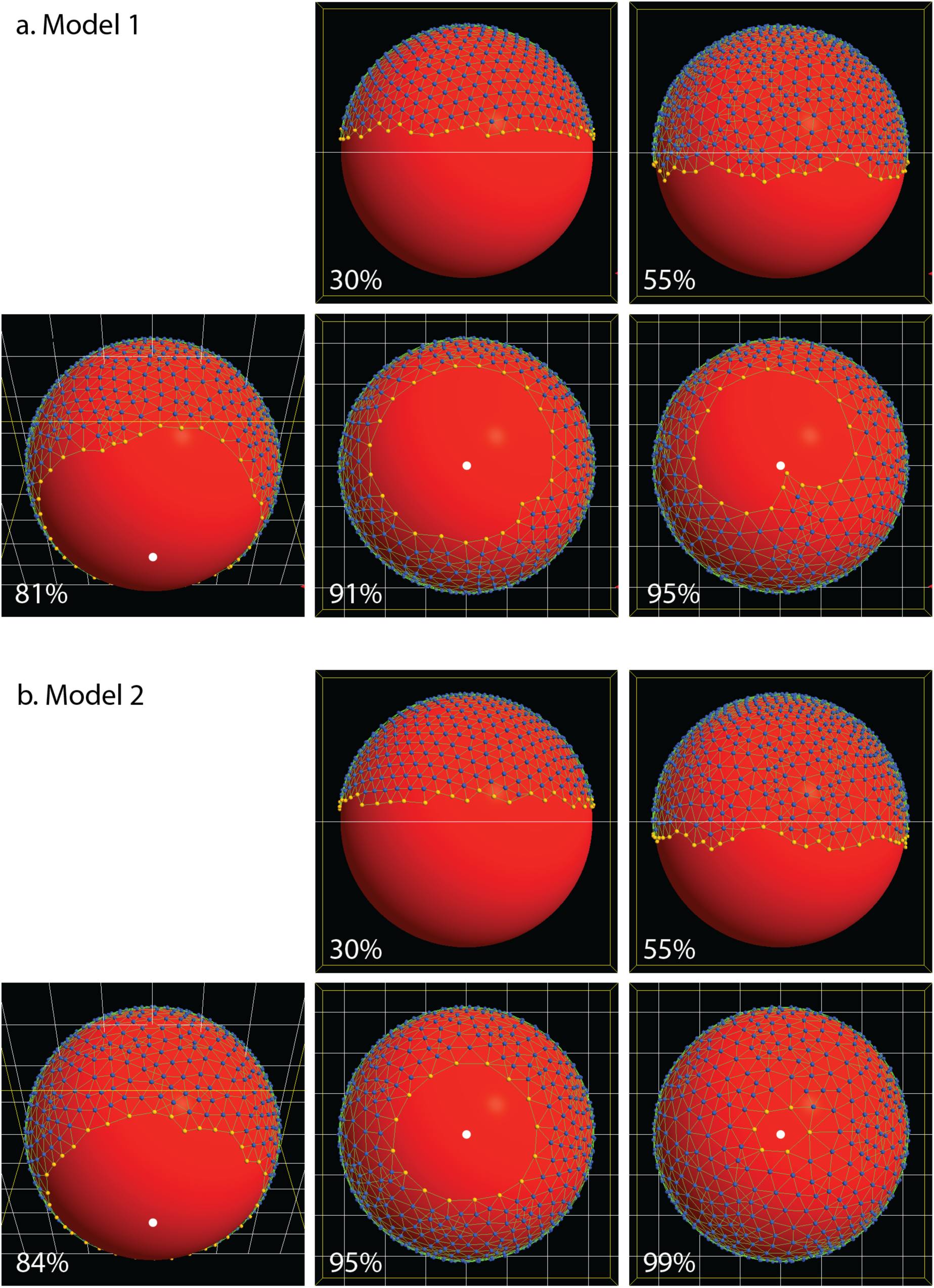
Epiboly progression in Model 1 and Model 2. Both of them (and except where noted, all subsequent examples) include dynamic bond remodeling, and cell division. Percentages indicate the embryonic stage (% epiboly). EVL cells are free to move between the leading edge and the internal region, but when a cell does so, it is recolored to match its new location, so although the cell populations change dynamically, the color pattern always demarcates the two populations. By 55% epiboly, all cell division has completed. Edge straightening: At 30% and 55% epiboly, the leading edge is ragged. By 90% epiboly, it straightens considerably in both models (though in Model 1, the straightness of the edge is lost later in epiboly). Tissue Forge has a programmable camera and we have it automatically begin rotating to a vegetal position when any point on the EVL margin reaches a mean polar angle *ϕ* = 0.75*π*. Vegetal pole is located at the bottom in lateral views (30%, 50% epiboly) and marked by a white dot otherwise (all later stages). a: Model 1, in which force is applied uniformly to all points on the leading edge. (See Movie S2.) Epiboly progress is synchronous through intermediate stages, but by around 90% epiboly, becomes visibly lopsided; the EVL leading edge is off-center relative to the vegetal pole, which is exactly in the center of the figure. By the end of the simulation at 95% epiboly, the edge is even more unbalanced, with a protrusion on the leading edge, toward the vegetal pole. The protrusion is typical in most runs of Model 1. Since all edge regions are not at a consistent polar angle, staging is scored by the mean polar angle of all the leading edge cells. b: Model 2, which adds force regulation. (See Movie S5.) The margin advances synchronously, remaining perfectly centered on the vegetal pole through the end of the simulation.

In Model 1 with or without cell division, vegetalward EVL expansion progresses quickly at first, and then slows down (Fig. 6a,b) as tension along the leading edge increases (Fig. 6c). Later, as the leading edge converges toward its point of closure, progress speeds up again, and tension drops (Fig. 6b,c). When cell division is included in the model, tension does not rise as quickly or as high. We expected that in the presence of cell division, tension might be kept low while cells are actively dividing, and start increasing more rapidly after cell division ceases, but Fig. 6c shows that in our model there is no marked difference in the rate of tension increase before and after cessation; the impact of early cell division is not limited to the period when cell division is occurring, but persists long past the time when division ceases (55% epiboly or *ϕ* = 0.53*π*; vertical line in Fig. 6c).

**Fig. 6.**
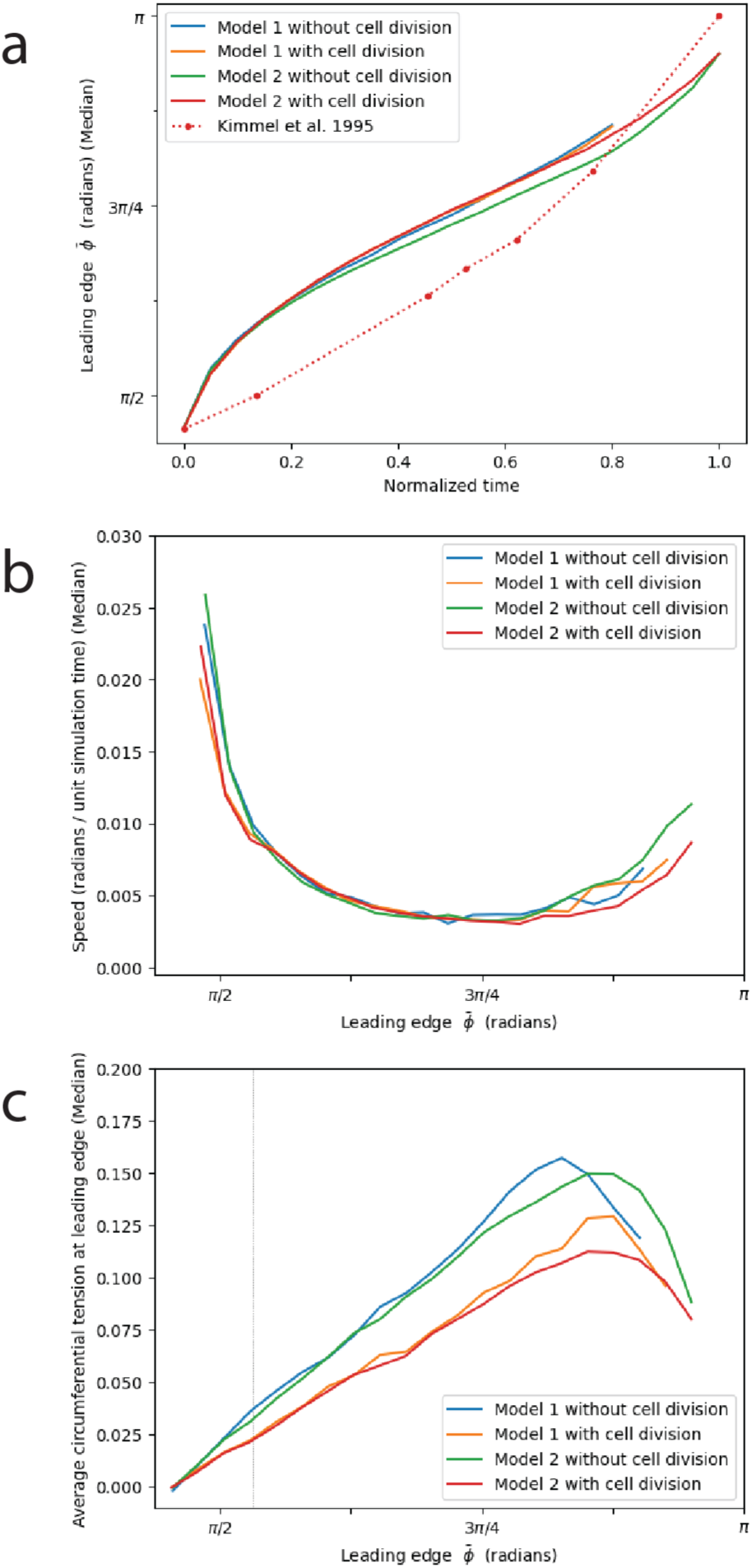
Measurements of simulation behavior over time. Consensus of multiple runs of Model 1 (with and without cell division, N=32 runs each) and Model 2 (N=46 runs each). a. Mean position of the EVL margin (polar angle *ϕ*), plotted against normalized time. (Time 0.0 and 1.0 represent, respectively, simulation start at 30% epiboly, and simulation termination, when the leading edge polar angle reaches 0.95*π*, or about 99% epiboly; see Methods.) The empirical data of Kimmel et al. [4] is superimposed over our simulated data for comparison (see Discussion). b. Speed of epiboly progression (plotted against the position of the leading edge on the horizontal axis; similar to plotting against embryonic stage; see Methods). c. Circumferential tension, measured along the leading edge. Vertical line marks the cessation of cell division (in those runs that had division enabled; orange, red) at 55% epiboly (polar angle *ϕ* ≅ 0.53*π*).

### Synchrony of epiboly progression and straightening of the EVL margin (Model 2)

In live embryos, the EVL leading edge progresses toward the vegetal pole faster on the dorsal side because of axial elongation [4]. But because our model considers only epiboly in isolation, without axial development, and is cylindrically symmetrical, we should expect epiboly to proceed symmetrically. The simple model presented above does not behave as expected; all points on the leading edge advance in approximate synchrony through early stages, but as epiboly progresses further, EVL advancement becomes increasingly unbalanced. The leading edge approaches the vegetal pole faster in some places than others (Fig. 5a, 91% - 95% epiboly; Movies S2-S4), creating a “lopsided” EVL. This imbalance likely arises from small local variations in force, due to stochastic effects or to the approximation in our force calculation (see Methods), leading to a mechanical positive feedback. As the most progressed region of the leading edge reaches the pole, it typically develops a small but distinct protrusion toward the pole (Fig. 5a, 95% epiboly; Movies S2-S4). To even out these imbalances and generate synchronous epiboly, we modeled a regulatory mechanism (Model 2), by modifying the rule for applying force to the different cells along the leading edge. In this regulated model, instead of applying a force of uniform magnitude all along the leading edge, we apply to each cell a force whose magnitude is proportional to its distance from the vegetal pole, so that lagging parts of the margin are pulled more strongly (see Methods for details).

In Model 2, the overall rate of tissue expansion, and the build-up and subsequent drop in tension along the leading edge, evolve over time exactly as in Model 1 (Fig. 6), but it now advances synchronously: all regions of the EVL leading edge advance at the same rate and converge simultaneously on the vegetal pole (Fig. 5b, Movie S5; Table 1, line 5). In Fig. 7a, we quantify the synchronization of leading edge advancement and compare the two models. The presence of lopsidedness in Model 1, and the possibility of enforcing synchrony by adding a regulatory process (Model 2), suggests that synchrony of an advancing edge in living morphogenetic systems such as zebrafish epiboly may not happen by default, but may instead require an active control mechanism, which to our knowledge has not been suggested in the empirical literature.

**Fig. 7.**
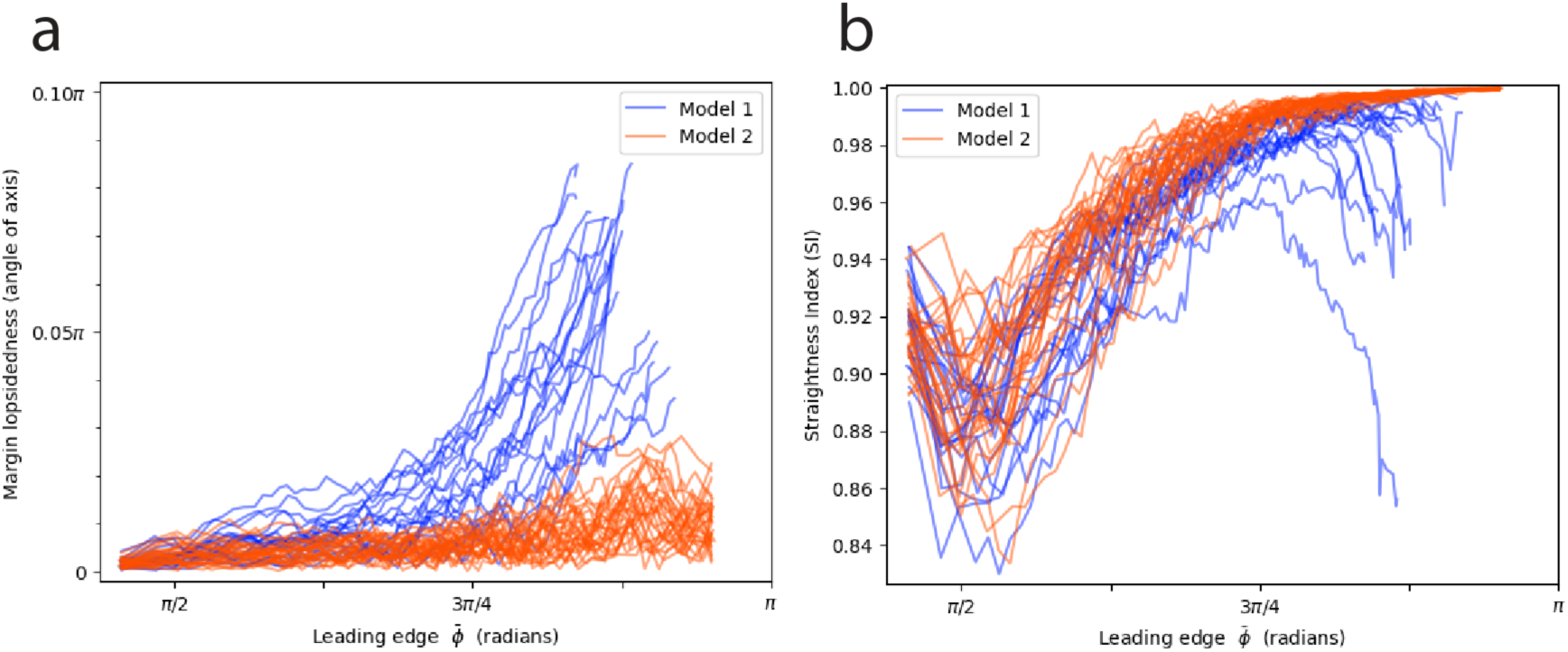
Regulation of the applied force is required for epiboly to proceed synchronously, but not for the straightening of the leading edge. Overlaid plots of individual simulation runs for Model 1 (N=20) and Model 2 (N=28). Margin lopsidedness (a) and Straightness Index (b) over time. See Methods for definition of the lopsidedness and Straightness Index metrics.

The presence or absence of such a regulatory mechanism may also affect the straightness of the leading edge over the shorter spatial scale of immediately neighboring cells. In living embryos, the EVL margin starts off loosely aligned, and becomes taut and straighter as epiboly proceeds [1]. We observe that the same behavior emerges in our model, so we quantified it by measuring the Straightness Index [19,20] or SI (see Methods for details) of the EVL leading edge over the course of epiboly in both models. The SI of both models (Fig. 7b) begins around 0.9, representing a somewhat loosely aligned arrangement resembling a living embryo (Fig. 5a,b, 30% epiboly). In both models, the margin then gradually gets straighter as epiboly proceeds (Fig. 5; Movies S2-S5; Table 1, line 5), recapitulating the behavior of live embryos, reflected in the rise of SI to around 0.99 in Model 1, and quite close to 1.0 (perfect straightness) in Model 2 (Fig. 7b). This is in striking contrast to the behavior of a purely elastic tissue, which becomes more and more jagged as it is stretched (Fig. 3b,c). At the end of epiboly in Model 1, SI frequently drops again, reflecting the protrusion which distorts the shape of the edge (Fig. 7b).

Our regulatory mechanism is therefore required for our model to achieve synchronized epiboly (only Model 2 proceeds in synchrony; Fig. 7a), but not for initial edge straightening (significant straightening of the edge occurs even in Model 1; Fig. 7b). We will return to a further investigation of the possible causes of edge straightening, after first discussing cell rearrangement, which bears on that question. Henceforth we further examine Model 2 only.

### Cell rearrangement and convergent extension

As the advancing EVL margin proceeds toward the vegetal pole, individual EVL cells migrate both into and out of the exposed edge (Fig. 8a,b). Net migration is out of the margin into the internal region, so the number of cells along the margin drops (Fig. 8a-c and Movies S2-S5). When cell division is added to the model, each division within the leading edge results in two leading edge daughter cells, adding to the population size of the leading edge (Fig. 8b); migration out of the leading edge increases to compensate (compare the “cumulative out” lines of Fig. 8a and Fig. 8b), so that the gradual net decrease in leading edge population size is the same (Fig. 8c, and the “Total margin cell count” lines of Fig. 8a and Fig. 8b). Cell rearrangement in our model is controlled by bond remodeling events: the making or breaking of bonds (adhesions) between cells. These events do not involve physical movement of the cells; each event is a change in connectivity, and movement follows, due to the resulting change in forces between cells. The topology of cell connections determines whether a cell is considered part of the margin; thus a cell is considered to have changed its identity from “margin” to “internal” or vice versa, and hence to have “migrated” in or out of the margin, when the change in the cell connection topology alters the cell’s exposure to the EVL edge. (Fig. 4b; see Methods for details.) In Fig. 8, we count these discrete events, rather than quantifying continuous cell movement.

**Fig. 8.**
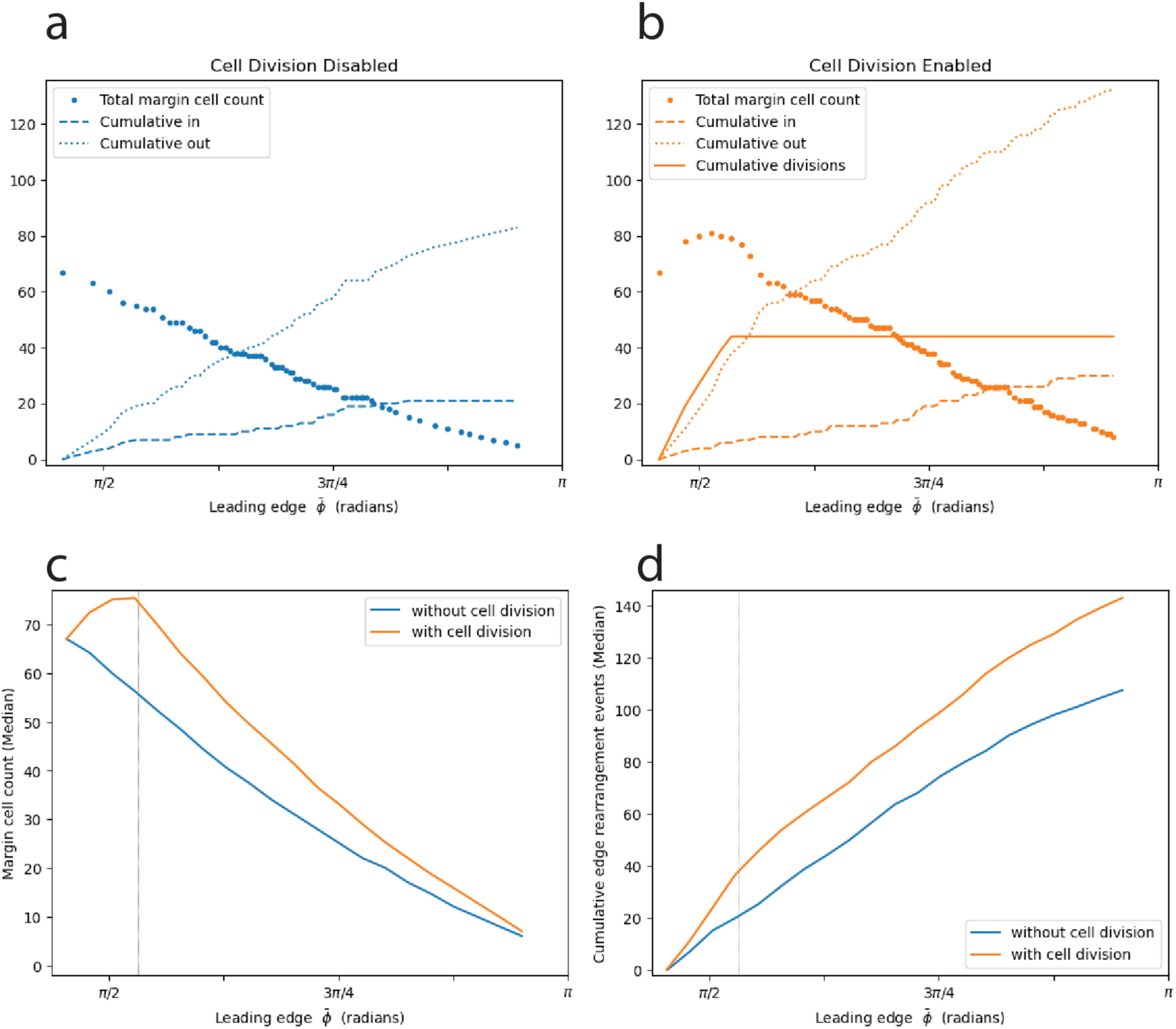
Cell rearrangement (Model 2). a, b: Cell rearrangement for representative examples of a single simulation run without cell division (a) and one with cell division (b), showing the changing number of margin cells over time, and its relationship to individual cell rearrangement events. Cumulative cell movements into, and out of, the leading edge are tracked, along with (in b) cumulative cell divisions.Leading edge cell divisions result in two leading edge daughter cells, and thus an increase in margin cell count. The change in margin cell count over time is the net result of these three types of events. Cell division ceases at 55% epiboly (polar angle *ϕ* ≅ 0.53*π*). c, d: Consensus of N=172 runs with and without cell division (86 runs each). Vertical lines mark the cessation of cell division (in those runs that had division enabled; orange). c: Margin cell count. d: Cumulative cell rearrangements at the margin (total amount of rearrangement: the sum of movements into, and out of, the margin).

To better visualize cell rearrangement, we performed virtual lineage tracing. Instead of color-coding particles dynamically according to the cell’s positional status (with cells changing color upon moving into or out of the margin, as in the earlier figures), we now color all particles the same (grey), except for a selected subset to label (red), and let each particle retain its initial labeled color throughout the simulation (Fig. 9a,b, Fig. S3, Movies S6-S8). At initialization (30% epiboly), we label either the EVL margin cells, or one tier of internal EVL cells, and track the movements of these cells within the EVL. When the leading edge cells are labeled (Fig. 9a, Movie S6), an enormous amount of cell rearrangement at the margin is evident. By the end of epiboly, the former margin cells have stratified into multiple tiers. In addition, the labeled and unlabeled populations become partially intermingled; some former margin cells are found quite far from the margin, and some internal cells are found much closer to the margin. When successively higher tiers of EVL cells are labeled (Fig. 9b, Fig. S3; Movies S7, S8), cell rearrangement and mixing are reduced, but some mixing is detectable even in tiers close to the animal pole, where cells move much shorter distances.

**Fig. 9.**
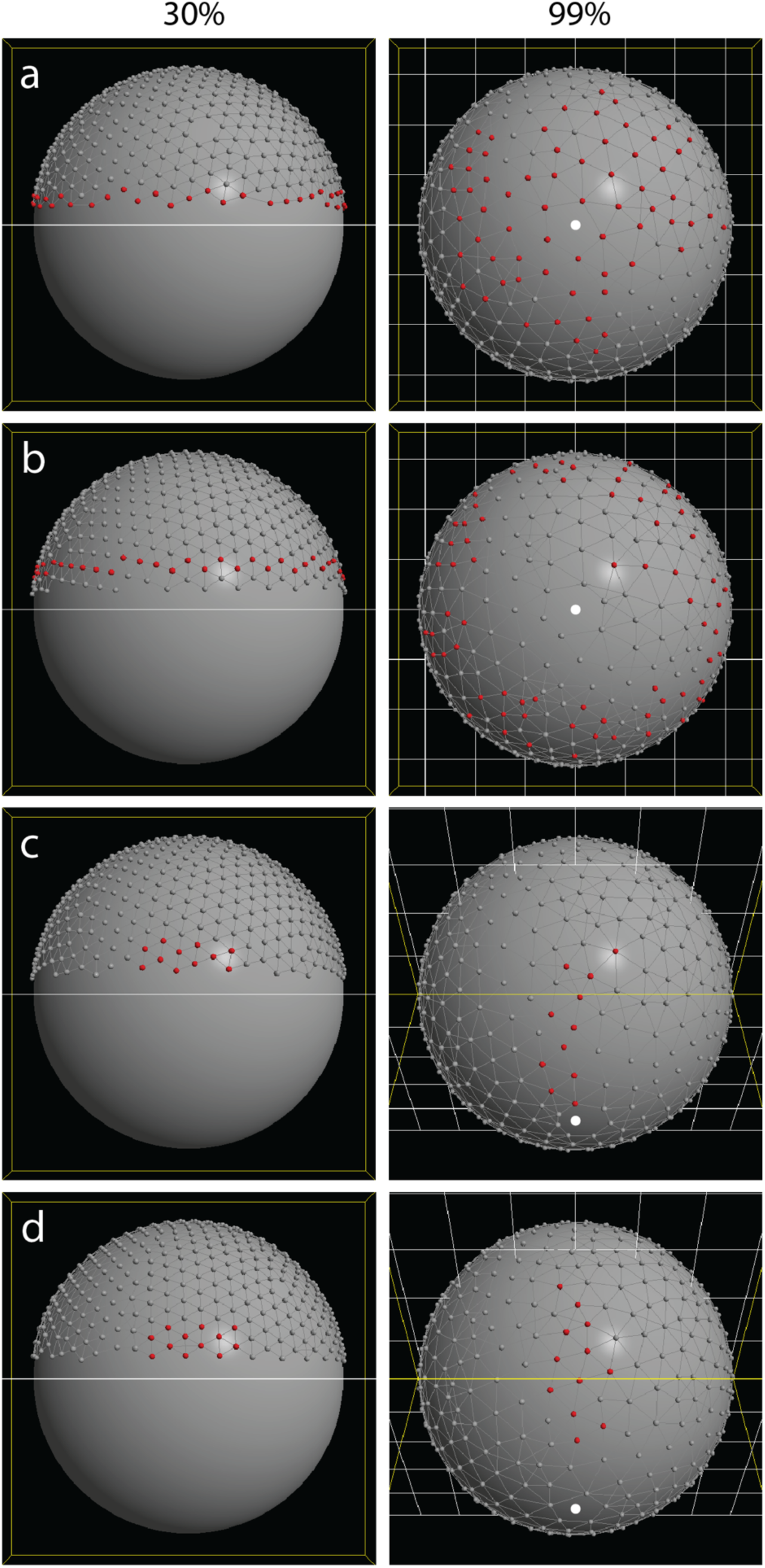
Lineage tracing in four simulation runs (Model 2). In each run, some cells were labeled red at initialization (30% epiboly, in lateral view, left); all cells retain their color through the entire simulation and are shown in their final positions (99% epiboly, right; with the vegetal pole marked by a white dot). Cell division was disabled so that any change in the geometry of the labeled region can be attributed simply to cell rearrangement, without any confounding effects of cell division. a. EVL leading edge cells are labeled. Many of these cells move away from the edge during epiboly, resulting in several tiers of labeled cells. By the end of epiboly, shown in vegetal view, a large area around the vegetal pole is composed mostly of cells originally at the leading edge. Intermingling of labeled and unlabeled cells is evident. (See Movie S6). b. Tier 1 of internal EVL cells are labeled. (See Movie S7). c. A patch of EVL cells on one side of the embryo is labeled, adjacent to the leading edge. In this example, the patch remains in contact with the leading edge, as is typical. Final view is from an oblique angle. (See Movie S9). d. A patch of EVL cells is labeled as in (c) and viewed from the same angles. In this case, the patch migrates away from the edge entirely. (See Movie S10).

To visualize convergent extension, we performed lineage tracing with patches of labeled cells, 30° wide and 15° high, adjacent to the EVL margin (Fig. 9c,d; Movies S9, S10). The longitudinal lengthening and circumferential narrowing of the patch are evident. (The patches also grow in surface area due to stretching of the individual cells, as can be seen by the increase in the distance between particles; thus, because stretch implies thinning in cells of constant volume, this is “convergence and extension and thinning”.) In some runs, the patch can be seen to widen a little bit as it stretches over the embryo equator, before narrowing to its final width as it approaches the vegetal pole. The final positions of the particles vary significantly between runs due to random mixing between the labeled and unlabeled regions, sometimes resulting in irregular patch shapes, or cells becoming isolated from the main patch; we selected examples with minimal mixing and more cohesive patches for Fig. 9c,d because it makes the change in patch proportions easier to interpret. Fig. 9c (Movie S9) shows an example in which the patch stays in contact with the leading edge, which is typical; Fig. 9d (Movie S10) illustrates that the patch can also migrate away from the edge entirely, in this case separated from the edge by two tiers of unlabeled cells by the end of epiboly.

### Straightening of the EVL edge is a robust outcome under parameter variation, and the rate of straightening is coupled to the rate of cell rearrangement

EVL edge straightening is emergent in our model; it was not explicitly designed in, but the model produces the phenomenon as in live embryos. Having shown that straightening takes place even in the absence of our regulatory mechanism, we sought to determine how it is generated in our model, by testing two other possible factors: a bias toward a straight edge inherent in the energy minimization criteria for bond remodeling; and circumferential tension in the leading edge. In each experiment we eliminated one potential contributor to the straightening, and examined the change in SI as epiboly progresses (Fig. 10, Fig. 11).

**Fig. 10.**
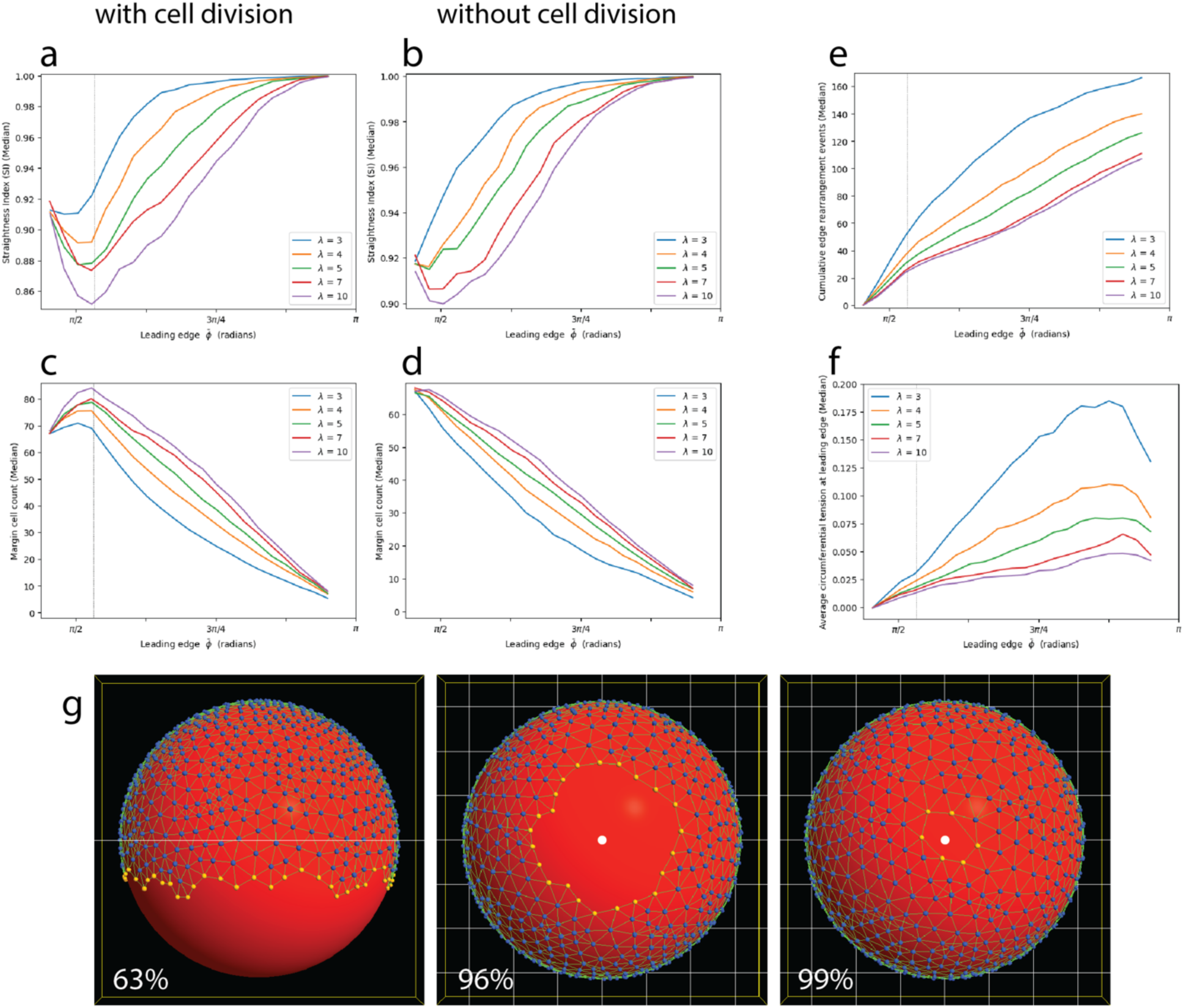
Rate of EVL edge straightening correlates negatively with the strength of the bond angle constraint. a-f. Change in several metrics over the course of epiboly. Each plot line represents median values of multiple simulation runs for each given strength (*λ*) of bond angle constraint (a, c, e, f: with cell division, 56 runs per treatment; b, d: without cell division, 32 runs per treatment), plotted against epiboly progress (the leading edge position over time). Vertical lines in a, c, e, f mark the cessation of cell division at 55% epiboly (polar angle *ϕ* ≅ 0.53*π*). a, b: Straightness index, in runs with cell division (a) and without cell division (b). The rate of increase of SI over the course of epiboly correlates negatively with *λ*. (The initial decrease in SI that occurs in most of the treatments in (a) likely is partly due to cell division at the margin, but not entirely: the same decrease occurs when cell division is disabled (b), though only at higher values of *λ*.) c, d: Margin cell count, in runs with cell division (c) and without cell division (d). The model allows cells both to enter and to leave the margin; the change in cell count over time is therefore the net change. Note that despite differences in the time course of margin cell population decrease, all treatments result in a similar net loss of ∼60 margin cells by the end of epiboly. The initial net increase in margin cell count that occurs in most of the treatments in (c) is due entirely to cell division during early epiboly, since no such increase occurs when cell division is omitted from the model (d). e: Cumulative margin cell rearrangement events. The total number of events correlates negatively with *λ*. f: Tension along the leading edge. The rate of increase in tension correlates negatively with *λ*. g: Selected time points in a single simulation run with *λ*=10. Straightening is delayed until much later in epiboly. See also Movie S11. Vegetal pole is located at the bottom in lateral view (63% epiboly) and marked by a white dot otherwise (96%, 99%).

**Fig. 11.**
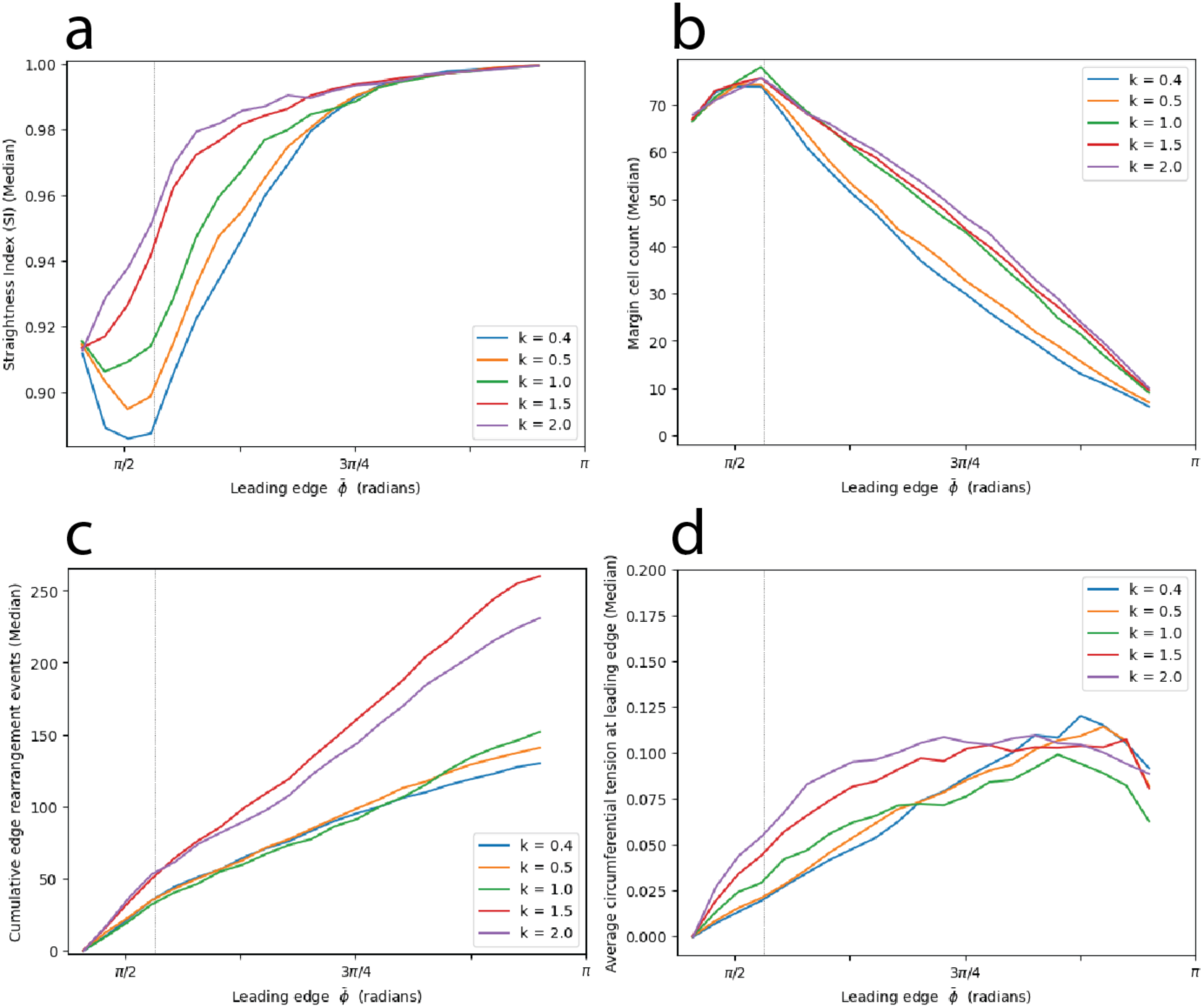
Rate of EVL edge straightening correlates positively with the spring constant of edge bonds (resistance of the edge to stretch). Spring constant was varied from our default value of 0.5. Vertical lines in each plot mark the cessation of cell division at 55% epiboly (polar angle *ϕ* ≅ 0.53*π*). Each plot line represents median values of 28 runs. a. Straightness index. b. Margin cell count. c. Cumulative margin cell rearrangement events. d. Tension along the leading edge.

Close examination of leading edge shape change by single-stepping through the simulation videos (Movies S2–S5) reveals that discrete straightening events are associated with discrete bond remodeling events at the margin, as an individual cell leaves the margin, and its two neighbors become bonded and close up the gap. This is a natural bias of the model, since the energy-minimization rules for bond remodeling favor a straight edge (see Methods). The strength of the constraint (the strength of the statistical bias toward the equilibrium configuration) is determined by the configurable parameter *λ*.

Bond remodeling events (removal of existing bonds or addition of new ones) result in changes to bond angles locally; the constraint acts by determining whether a candidate event will be accepted or rejected, according to the energy minimization criterion (see Methods for details). The stronger the constraint (the greater its *λ*), the more likely it is for unfavorable events (those causing the configuration to move further from the favored one) to be rejected. Thus we expect this algorithm to keep bond angles near their target value. The constraint is applied throughout the EVL; internally to the EVL, we configure the constraint to favor 60° angles, a hexagonal packing. (Indeed, this is why we implemented the constraint, as it enables the tissue to stretch without developing holes.) Along the edge, the target value is a straight boundary (near 180°, with an adjustment for the curved embryo surface). To test whether the straightening of the EVL edge is a consequence of this built-in bias, we weakened the constraint along the EVL edge by reducing *λ* in that region. We modified *λ* specifically for candidate events that add or remove a bond between two margin cells. For all other bonds (those involving at least one non-margin cell), *λ* was held constant. Reducing *λ* should lead to more frequent acceptance of candidate events that increase the deviation of the bond angles from their target value near the edge, interfering with the ability of the edge to straighten. Moreover, during each such event at the margin, a cell enters or leaves the margin, so reducing *λ* should lead to a greater number of such rearrangements, which we can measure.

However, our results contradicted this expectation (Fig. 10). Reducing *λ* from our default value of 4.0 actually caused straightening to occur faster, completing earlier in the progression of epiboly (Fig. 10a,b). In contrast to the effects on straightening, cell rearrangement at the edge did behave as expected, becoming more active. The net change in the number of margin cells between the beginning and the end of epiboly was about the same regardless of treatment, but it occurred earlier in the simulation as *λ* decreased (Fig. 10c,d); and the total number of cell rearrangement events, with cells moving both in and out of the margin, increased (Fig. 10e), as did the circumferential tension, measured along the leading edge (Fig. 10f).

We therefore tested the effect of increasing the value of *λ*. This had the opposite effect: the total amount of cell rearrangement decreased (Fig. 10e), straightening took place more slowly (Fig. 10a,b), and tension along the leading edge increased more slowly (Fig. 10f). Morphologically, the ragged edge can still be observed late in the simulation, finally straightening after a delay (Fig. 10g), and when viewed dynamically (Movie S11), the edge appears rigid and resistant to deformation. The strength of the constraint is therefore negatively correlated with both the overall amount of neighbor exchange and the rate of edge straightening. This is counterintuitive: the higher the *λ*, the stricter the constraint, i.e. the more strongly and quickly the configuration should converge on its target value (a straight edge). Instead, we find that a stronger constraint does not encourage edge straightening, but slows it down. We may understand this result as follows: fluctuation is an integral component of a stochastic, energy-minimization algorithm. Without fluctuation, a system may get stuck in a local energy minimum, and not be able to reach a more global minimum (the target configuration: a straight edge). Fluctuations (temporary excursions in the “wrong” direction, away from the lowest energy state) allow quicker discovery of a path to the global minimum. Thus the higher the *λ*, the more often unfavorable rearrangements will be rejected, leading to a slower rate of cell rearrangement overall; and thus to slower convergence on the target configuration. Stated differently, increasing *λ* leads to a greater potential energy difference between configuration states (greater favorability of a straight edge), but also raises the activation energy of the transformation (larger fluctuations required to get there). The situation is analogous to a thermodynamically favorable, but kinetically slow biochemical reaction such as the breakdown of glucose: it releases energy because it is favorable, but it requires an input of ATP and catalyzing enzymes, to overcome the activation energy.

More intuitive is the relationship between straightening and tension along the edge. The rate at which tension increases as epiboly proceeds (Fig. 10f), and the rate of edge straightening (Fig. 10a,b), both correlate negatively with *λ*. Increasing tension may pull the edge taut and straighten it. We attempted to test this more directly by manipulating the spring constant of the bonds joining pairs of margin cells (representing the stiffness of the bonds or their resistance to deformation), because the tension in a bond is the product of its spring constant and its deformation (Fig. 11). We found that the rate of edge straightening correlates positively with the spring constant (Fig. 11a). The effects of the spring constant on cell migration, and on the tension along the leading edge, were less consistent (Fig. 11b-d). Treatments leading to faster straightening (i.e. higher spring constants, Fig. 11a) appear to generate, above a certain threshold, more cell rearrangement events yet slower net change in total edge population, but not in a simple pattern (Fig. 11b,c).

### In silico “laser cut” experiments demonstrate viscoelastic deformation of the model EVL

Our epiboly model displays viscoelastic deformation; internal remodeling of cell adhesions allows for cell rearrangement, greatest near the EVL leading edge late in epiboly, associated with relaxation of tension (Fig. 6c). Although individual bonds are perfectly elastic, this viscoelastic behavior of our model at the tissue level allows the EVL to adopt its new spherical shape permanently. To demonstrate this, we performed the following experiment. When the leading edge has progressed to a mean polar angle of 0.9*π* (∼97.5% epiboly), we disable the exogenously applied forces (similar to a laser cut experiment on a live embryo, disrupting the EVL-yolk junctions through which exogenous forces are transmitted), and allow the simulation to continue running without them. To test the instantaneous recoil reaction of the cell sheet, we simultaneously disable the active bond remodeling; thus the topology of the connected cell network is fixed. Under these conditions, a completely elastic material would return to its original shape (Fig. 3d, Movie S1). In contrast, a viscoelastic material under tension would recoil a small amount, reflecting its short-time-scale elasticity, but largely maintain its overall shape, reflecting its long-time-scale viscosity. This is indeed what we see (Fig. 12, Movie S12). Most of the cell sheet stays in place, except for a small area near the margin, which recoils a short distance and then quickly stabilizes (plateau in Fig. 12). At this point the tissue is still under tension (all bonds are stretched beyond their resting length), but the net forces on each cell are balanced because of the reorganized configuration. Although exogenous vegetalward forces have been removed, the tissue’s own constriction along the leading edge gives rise to an internally generated vegetalward force due to the edge being located below the equator. Initially the edge begins to recoil because internal animalward forces outweigh these vegetalward ones. As the edge recoils toward its new equilibrium, the bonds along the leading edge become stretched, causing the vegetalward force to increase until it balances the animalward force, and recoil comes to a halt. In a second phase of the experiment, we then re-enable bond remodeling, making the configuration more labile, and the cell sheet undergoes an additional large contraction (Fig. 12, Movie S12), further demonstrating that the stability of the tissue even under tension arises from its internal organization.

**Fig. 12.**
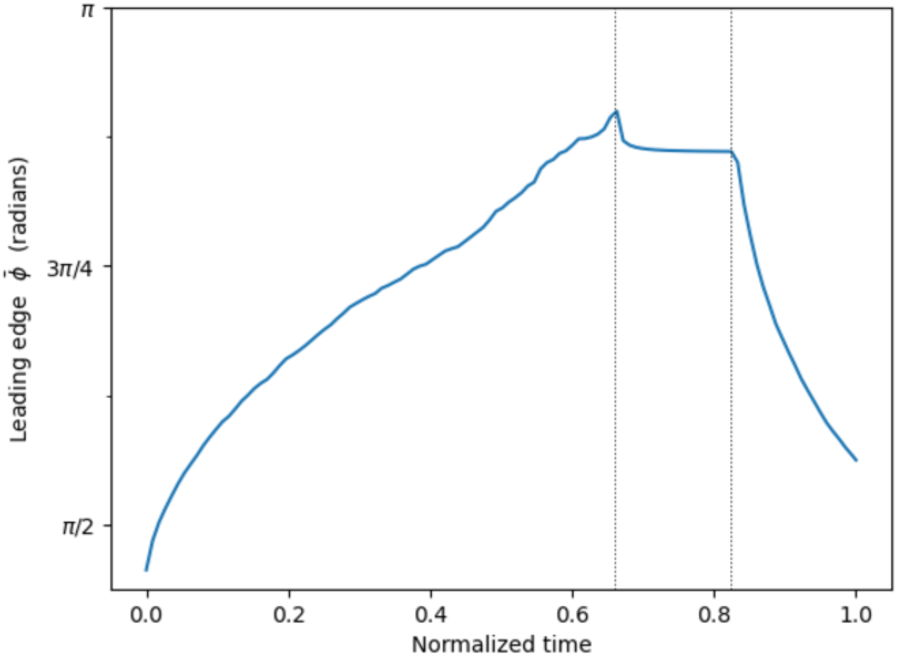
Response of the reorganized model EVL to experimental disruption of forces demonstrates its viscoelasticity. Mean polar angle of the EVL margin in a single representative example, plotted against normalized time; vertical marker lines indicate the start of the first and second phases of the experiment. When the leading edge reaches a polar angle of 0.9*π* (∼97.5% epiboly; first vertical line), we disable both the exogenously applied force and the internal bond remodeling. The reorganized EVL undergoes minimal instantaneous recoil (short term response) and stabilizes. But the tissue is still under tension, as demonstrated by the second phase of the experiment (to the right of second line): we re-enable bond remodeling, allowing continued cell rearrangement to occur, resulting in further recoil (long term response). See Movie S12.

## Discussion

We have formulated a simplified, center-based model of zebrafish EVL morphogenesis during epiboly, using the Tissue Forge modeling framework. EVL expansion and shape change in the living embryo requires not only the deployment of cell behaviors and mechanical forces capable of generating the needed transformations, but also the ability to withstand and respond to those forces while maintaining mechanical tissue integrity. Building our epiboly model was a discovery process that provided insight into the challenges faced by living embryos.

Our model is an extremely simplified representation of the embryo; it contains no explicit representation of cell boundaries, nor of EVL cell thickness, nor of the deformable yolk; and it considers epiboly independently of axial development. Yet, this model captures a number of the features of EVL epithelial expansion and shape change. First and foremost, its objects, structure, and behavioral dynamics result in a coherent sheet of particles which, like an epithelium, can be stretched by an applied exogenous force and will tear if over-stretched; but given an optimal choice of parameters, the tissue can respond to that force by stretching without tearing. The bond-remodeling enables the tissue to undergo internal viscoelastic rearrangement, transforming from a spherical cap to a complete sphere as it engulfs the yolk, without buckling.

### Synchrony of epiboly progression

Our simpler model, Model 1, gradually becomes lopsided over the course of epiboly; different regions of the leading edge extend toward the vegetal pole (the point toward which the applied forces are all directed) at different rates. Live embryos have a dorsal-ventral axis, and axial elongation results in the dorsal side extending faster; in fact, it expands beyond the vegetal pole, converging with the rest of the leading edge on the ventral side [4]. Live EVL cells therefore don’t have the pole as their destination; and besides, they are not driven by long-distance goal directedness, but by local forces. Moreover, the dorsal-ventral asymmetry in epibolic extension is consistent and predictable; it is in this sense that we can refer to epiboly in live embryos, although asymmetric, as synchronous: all sides converge at their common (slightly ventral) destination simultaneously. In contrast, our model embryo does not have a dorsal-ventral axis, and its destination is explicitly defined as the vegetal pole, so the lopsidedness of Model 1 is quite a different behavior: it does not converge on its destination in a predictable way, and the point that reaches the vegetal pole first can be located at any position along the circumference of the leading edge. An additional mechanism was required for the model to capture the same synchronous behavior exhibited by live embryos, suggesting that such a mechanism may also be required in living embryos, to coordinate their synchronous behavior.

Our improved model, Model 2, regulates its EVL boundary progression so that it advances synchronously. We enforced synchrony of the leading edge by applying a simple regulatory control to the pulling force of epiboly, based on the distance of each particle from the vegetal pole. In other words, we supposed for the purposes of calculation, that a force-generating mechanism might be sensitive to the polar position of the cells. We do not mean to suggest a direct mechanistic explanation of force regulation, but we can ask whether the general principle of such regulation is sensible.

We are not aware of any study addressing the regulation of synchrony during epiboly. Do zebrafish have such a regulatory mechanism? We suggest that the synchrony of epiboly progression may offer insight into the potential roles of different proposed force generation mechanisms in epiboly. We can ask, theoretically and experimentally, whether each proposed mechanism has the properties necessary to explain the synchronous behavior.

For example, the cable constriction model of epiboly proposes that epiboly is driven by constriction of the actin band surrounding the embryo in the external part of the YSL (the e-YSL). This hypothesis seems poorly suited to explain the synchrony of epiboly. At any given point during epiboly, the actin band approximates a circle, whose center lies on the embryo axis. The net constricting force acting on any given point along the edge will be a vector pointing toward the center of that circle. At the equator, this force vector would be perpendicular to the direction of EVL expansion; and as epiboly progresses, that force vector and the direction of EVL expansion come into greater and greater alignment. Therefore circumferential constriction would be more efficient closer to the pole, so it should pull leading regions more strongly, and lagging regions more weakly, increasing lopsidedness rather than resolving it.

In contrast, other proposals hold that epiboly is driven by forces within the e-YSL directed in an animal-vegetal direction, for example by endocytosis [6–10] or by tension in actin filaments aligned perpendicular to the band rather than along it [11]. In these scenarios, the generated force may be proportional to the thickness of the e-YSL (the distance between the EVL leading edge and the vegetal limit of the e-YSL); and if the e-YSL itself has a variable thickness, then a region of EVL adjacent to a narrower force-generating region of e-YSL would experience less force, thereby providing a regulatory mechanism quite similar to what we have modeled (Fig. 13).

**Fig. 13.**
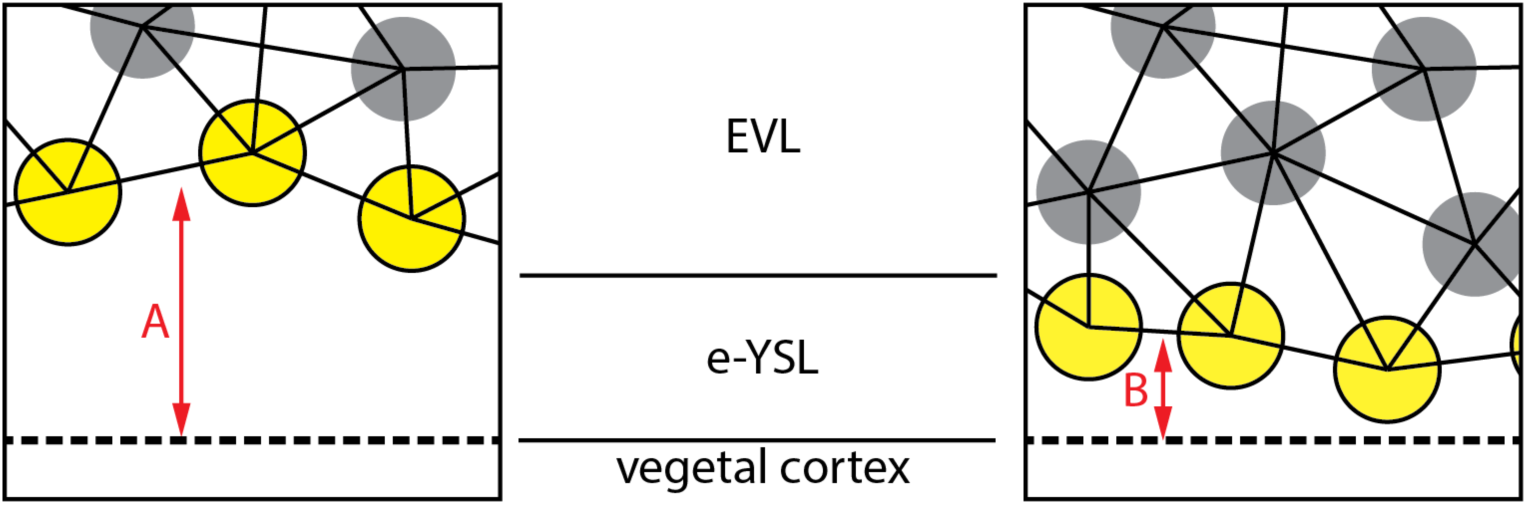
Hypothetical dependence of the magnitude of applied force on the thickness of the e-YSL. Yellow particles represent the cells of the EVL leading edge, in separate regions of the embryo, one lagging (A) and one leading (B). The e-YSL is the region between the EVL edge animally, and the main yolk cortex vegetally. If the boundary between the e-YSL and vegetal cortex is itself advancing synchronously; and if the pulling force is generated by cumulative processes distributed within the e-YSL (e.g. endocytosis), then the magnitude of the force applied to the EVL in each region should scale with the available surface area of the nearby e-YSL, i.e. with its thickness (distances A and B). This could lead to stronger forces exerted on lagging regions than on leading regions, and thus a negative feedback keeping all regions of the leading edge advancing in synchrony.

### Cell rearrangement, EVL edge straightening, and tissue deformation

For the EVL to accommodate the tissue shape changes of epiboly, cell rearrangement is required. Extensive rearrangement has been documented at the leading edge [1,15], resulting in a decrease in the number of EVL margin cells as epiboly proceeds. We would expect significant rearrangement in all regions of the EVL below the equatorial zone, because there, epiboly is a process of convergent extension: to extend toward the vegetal pole, the tissue narrows circumferentially, while lengthening longitudinally. Above the equator, the situation is different: there is no convergence, and the tissue extends both circumferentially and longitudinally, as it stretches over the widest part of the embryo. The equator does not mark a sharp disjunction between these two deformation patterns, but a transition zone, where tissue expansion approximates a cylinder stretching along its length, with the circumference of the cylinder not changing significantly. The character of deformation therefore transforms smoothly and gradually between the animal and vegetal poles; and cells that cross the equator during epiboly progress through all of these deformation regimens. In our model, cell rearrangement takes place throughout epiboly, and in all regions of the EVL (Fig. 8, Fig. 9, Fig. S3, Movies S6-S10), greatest near the leading edge. Because cells near the animal pole are expected to move only short distances during epiboly, it might be expected that little rearrangement would occur there. But in our model as revealed by lineage-tracing experiments, mixing of labeled and unlabeled populations takes place even near the animal pole, albeit to a much lesser degree. Empirical observations in living embryos reveal similar large rearrangements, at least at the EVL margin where these studies have focused their attention, and point out that this can occur without impairing the important permeability barrier function of this epithelium [1,15].

We considered the potential roles of three possible algorithmic features of our model, in bringing about the straightening of the EVL edge. First, the regulation of the pulling force (not included in Model 1, Fig. 5a, Movie S2, but added in Model 2, Fig. 5b, Movie S5) might promote straightening of the EVL edge by adjusting the relative force on nearby cells, in the same way that it prevents lopsidedness by adjusting forces over longer spatial scales. Second, the bond angle constraint of our bond remodeling rules, with its inherent preference for certain cell packing geometries (Fig. 10), might create a statistical bias in favor of straightening. And finally, circumferential contraction due to tension in the bonds between leading edge cells (Fig. 11) might pull the edge taut. We found that the rate of straightening does not depend on the regulation of the pulling force; and that the bond angle constraint, although in principle introducing an energetic bias in favor of straightening, in fact slows straightening down. This correlation, the reverse of what was expected, may be mediated by indirect factors: a loosened constraint (lower *λ*) leads to a steeper increase in tension along the EVL edge, and to a greater total number of cell rearrangement events (although the total net migration out of the edge remains roughly constant), and either of these factors might promote straightening. In all of these experiments, we were not able to prevent EVL straightening, but only to modulate the rate at which it occurs. This suggests that straightening is a robust emergent property of our model, probably controlled redundantly by a complex combination of factors (similarly to how control mechanisms tend to work in living systems).

How can these experiments help us understand EVL edge straightening in living embryos? Some of the features of our model correspond directly to living features. Bonds represent adhesions, and their properties (resting length and spring constant) relate to properties of cell adhesion such as adhesion strength, and the stiffness or elasticity of adhering cells. The pulling force, though defined generically and independently of any particular force generation model, relates to both compressive and contractile forces in embryos that are likely propagated along structural elements like cytoskeleton and cell junctional complexes, and driven by cytoskeleton-associated motor proteins. In contrast, the bond angle constraint – a key player in these experiments – is not a literal representation of an easily identifiable cell structure or process. We introduced it of necessity, to enable our model tissue to respond to large forces without disintegrating, thereby capturing the ability of living tissues to tolerate large deformations and to remodel themselves. At the level of the local cell neighborhood, the constraint describes a tendency toward a hexagonal packing, and thus embodies the tendency of cells in a living epithelium to pack efficiently, a consequence of a complex interplay between surface tension, cell adhesion, the structure and dynamics of the cytoskeleton and extracellular matrix, and mechanosensitive signal transduction (reviewed in [21]). That relaxing the constraint leads to more rapid cell movement both into and out of the margin (Fig. 10e), along with an acceleration of EVL edge straightening (Fig. 10a,b), suggests that rapid cell neighbor exchange may promote a transition of tissue near the leading edge from jammed (solid-like) to unjammed (fluid-like) behavior, as described in [21,22] for the expansion of the serosa in embryos of the beetle *Tribolium castaneum*, an epithelial engulfment of the embryo strikingly similar to zebrafish epiboly. There, irregular cell packing and rapid neighbor exchange near the serosa leading edge leads to increased cell rearrangement and circumferential shrinkage, required for tissue deformation and engulfment. The same dynamics appears to be in play in our model, underpinning both the deformation required for engulfment, and the straightening of the leading edge. The question then arises, whether the same can be shown in live zebrafish embryos: does rapid rearrangement represent a fluidization of the tissue, required for edge straightening and perhaps for the overall tissue shape change?

Our model allows all available topological transformations during bond modeling, including the movement of cells either into, or out of the leading edge. We make no a priori restriction, but allow the underlying physics to determine the dynamics. Experimental studies have documented only movement out of the leading edge [1,15]. In our model, the cells do occasionally move into the leading edge, but with a large net outward migration. Our plot of the margin population size in simulations when cell division is included (Fig. 8c) is strikingly similar to the equivalent plot of the same measurement in live embryos [15]. Whether individual cells occasionally enter the leading edge in live embryos is an open question; answering it may have implications for understanding the mechanisms controlling cell rearrangement.

A drawback of our model is that it contains no explicit description of cell shape. We have emphasized that cell shape change and rearrangement are complementary components of tissue shape change; but they can also act to enhance one another. Studies in zebrafish have described the elongation of EVL margin cells as they undergo rearrangement [15] and have shown that cell shape change, like tissue extension, depends on actomyosin [1]; and cell shape is thought to have a critical role in potentiating the fluidization of tissues and thereby enabling cell rearrangement [21,22].

### Viscoelastic deformation

In viscoelastic deformation, internal rearrangements are elastic (reversible) over short time scales, but become permanent over longer time scales, as internal relaxation reduces stress. In our model the elasticity comes from the individual bonds, and the internal relaxation comes from the remodeling of the bonded interactions (topological restructuring of the network of connected cells), which results in bonds under greater strain being replaced by bonds under less strain. In living embryos, internal structure is of course much more complex; but cell division and cell rearrangement contribute to internal relaxation [14]. When the exogenous force is removed from our model near the end of epiboly, while suppressing bond remodeling, the EVL undergoes only a small amount of recoil (Fig. 12, Movie S12). This represents a short-term response to the removal of the force, and indicates that the overall deformation of the EVL is stable. When we then re-enable bond remodeling, representing a longer-term response, the model EVL rapidly contracts, demonstrating that the tissue is still under tension, and can reverse its shape change by reversing the remodeling of the bonds. That such reversal has not been observed in real embryos allows us to predict that cell junction remodeling and cell rearrangement act as a ratchet: their reversal is likely prevented, making the change in tissue shape permanent. EVL expansion during epiboly is a passive response to external forces, and it is possible that the cell rearrangements within the EVL required to accommodate that expansion are likewise a passive response to those forces; but it has been suggested that there may be some active component to the rearrangement [1,14,15]. EVL cells may actively increase their junctional remodeling activity to promote (or at least allow) rearrangement during epiboly and then decrease it to reinforce their final configuration, or they may even actively regulate the direction of cell rearrangement by favoring certain junctional alterations over others in a location-dependent manner, creating the ratchet.

### Time course of epiboly

Kimmel et al. [4] plotted epiboly progression (% epiboly) against developmental time and showed it to be linear (except for the pause at germ ring and shield stages, during which the position of the EVL margin holds steady at 50%). For comparison with our results, we superimposed their data on ours in Fig. 6a (omitting the stationary pause so as to compare apples to apples), thus replotting it in terms of polar angle *ϕ* instead of % epiboly. Polar angle produces a better representation of the speed of leading edge advancement, because the arc length traversed by a point on the edge is directly proportional to the distance traveled, while % epiboly is not. Viewing the Kimmel et al. data in this way reveals a speed-up in the later stages of epiboly, just as in our simulations. In contrast, the simulations do not match the empirical data as well during the early and middle parts of epiboly; during these earlier stages, living EVL appears to advance at a fairly constant speed, whereas our simulated EVL starts out progressing quickly, then rapidly decelerates (Fig. 6a,b, Movies S2-S5). The final speed-up suggests that tissue expansion (both living and simulated) meets with less resistance as the leading edge approaches the vegetal pole, likely a consequence of the geometry: the deformation at the leading edge, when near the equator, approximates a stretching cylinder, whereas near the vegetal pole, it approximates a shrinking hole in a flat plane, similar to wound healing. Our method of applying force (keeping the force per unit length of margin constant over time) is simple but arbitrary, and the differences between the living and simulated data suggest that our method may embody incorrect assumptions; live embryos may very well modulate the force in a more complex manner, and/or modulate the internal tension of the EVL, to control the time course of epiboly.

The EVL in our model comes under greater tension as it stretches; and tension along the leading edge then drops again as the leading edge shrinks, late in epiboly (Fig. 6c). These changes in tension over time may be related to the observed changes in speed of epiboly, since epiboly initially slows during the period of increasing tension, and speeds up again as tension subsequently drops; however, the relationship between the two must be indirect, since the addition of cell division to the model leads to a reduction in tension build-up but little change in the speed of epiboly (Fig. 6b,c). The reduced tension when cell division is present is consistent with experiments in live embryos [14] showing that anisotropic tension in the EVL leads to oriented cell division, which in turn reduces tension anisotropy, a negative feedback loop that keeps tension low, promotes epibolic expansion, and helps maintain tissue integrity. Cell division in our simple model is oriented randomly, so cannot have an anisotropic effect directly; however, if forces within the EVL are anisotropic under conditions of stretching, then even randomly oriented daughter cells would likely be pulled into alignment by those forces after division. This is supported by the fact that cell division leads to more cell rearrangement (Fig. 8). The effect of cell division on tension appears not to be directly dependent on ongoing cell division, but is felt consistently throughout epiboly, even after division has ceased (Fig. 6c), suggesting that reduced tension results simply from the presence of greater numbers of smaller cells, rather than from the division event itself. We plan to investigate the role of anisotropic forces and oriented division further in future work.

## Methods

We use the toolkit of mechanical components provided by Tissue Forge [18] to represent the epiboly-stage zebrafish embryo as follows.

### Model objects

#### Particles

Tissue Forge particles are point masses, and therefore each particle represents the center of mass of an object; it is up to the simulation code that uses Tissue Forge, to define the nature of that object and its scale, from the atomic to the organismal. We use a single large particle to represent a composite of the yolk cell and deep cells; and otherwise we use one much smaller particle for the center of mass of each EVL cell. Tissue Forge particles are spherical, with a defined *particle radius*, which affects only the graphical rendering, and not the actual physics of particle interactions. The spherical surfaces of our EVL particles therefore don’t represent cell surfaces, nor do the particle radii represent cell size. Instead, we define a *cell* represented as a particle combined with an assigned *cell radius* distinct from (and larger than) the particle radius that Tissue Forge tracks (Fig. 2). (The foregoing does not apply to the yolk particle, which is treated more simply; its particle radius represents the radius of the living yolk cell plus deep cells.) Since EVL cells are squamous epithelial cells, their representations in our model are conceptually flat rather than spherical, and our cell radius applies only to the extent of the flat cell within the sheet – i.e. the radius of the apical or basal cell surface – not to the cell’s thickness. The EVL particle radius determines the thickness of our EVL layer, but we do not attempt to model the changing thickness of living EVL (apical-basal cell height) as it spreads. Since living epithelia are seamlessly connected with no gaps, we consider all the space within the cell sheet to be part of the cells, which therefore represent irregular shapes, though we treat them as idealized disks. We do not explicitly represent those irregular shapes. The potentials assigned to interacting particles (described below) determine the repulsion between them due to volume exclusion (i.e., resistance to compression) and thus the distance limit of approach; and are based on the cell radius rather than the particle radius. The model cells are therefore, like live cells, deformable (their shapes can change) but incompressible (their volumes cannot). Any EVL cell elongation (elongation of the apical surface) is implicit in the arrangement of particles, since any anisometry of center-to-center distances can be interpreted as an anisometry of cell shape. Actual cell size can likewise be inferred from those center-to-center distances; the assigned cell radius represents a target or equilibrium radius (and therefore apical surface area), from which the actual surface area may diverge depending on the cell’s mechanical interactions with its neighbors. (E.g. a cell whose area is greater than the target can be interpreted as being stretched, and, because the cell volume is constant, thinned to compensate.)

In addition to radius, Tissue Forge particles also have a second property, “mass”. In the context of our simulation, the mass property is better thought of as the drag coefficient. Tissue Forge supports two modes of particle dynamics: Newtonian and overdamped. Newtonian dynamics is represented by the familiar F=ma, and is appropriate for planets, billiard balls, and molecule-to-molecule interactions; overdamped dynamics models fluid environments, where inertial forces are negligible compared to fluid drag, which dominates. We use overdamped dynamics, in which force is proportional to particle velocity rather than to acceleration; the constant of proportionality thus represents the drag coefficient.

Tissue Forge particles are instantiated as members of user-defined particle types, which are classes of particles with predefined mass and particle radius. We define three particle types for the simulation: the yolk cell is its own type, and we define both an “EVL margin” and “EVL internal” type to distinguish EVL cells positioned at the EVL margin (the cells which, in live embryos, are bound to the yolk by tight junctions and subjected to exogenous force generated in the yolk and propagated via that mechanical coupling) from the rest of the EVL cells (which in the embryo are not bound directly to the yolk and are separated from it by the deep cells). While the Tissue Forge notion of “particle type” can be used as an analog to biological cell type, in our case the two EVL particle types are considered the same cell type, with similar mechanical properties (typically identical properties, though in certain experiments we vary their parameters separately). Tissue Forge particles can be dynamically reassigned to a new particle type, and our EVL cells switch freely between “EVL internal” and “EVL margin” during cell rearrangement whenever conditions lead to a margin cell moving inward from the margin, or an internal EVL cell moving outward to the margin. This particle-type distinction facilitates the distinct manipulation and analysis of the two subpopulations, including subjecting them to different environmental effects (e.g. distinct forces); specifying different behaviors (e.g. distinct particle connection topologies); taking separate measurements (e.g. plotting tension measurements between margin particles only); and applying separate user interface properties (e.g. displaying each subpopulation in its own color).

### Model processes

#### Potentials and bonds

Tissue Forge potentials are functions defining the energy (and hence the attractive and/or repulsive force) between two particles as a function of the distance between them. Tissue Forge supports a variety of potential functions common in physics modeling; we use a simple harmonic potential throughout, which behaves exactly as an ideal spring, defined by a resting length and a spring constant. We set the resting length equal to the sum of the cell radii of the two interacting particles/cells, such that at equilibrium, the invisible but inferred surfaces of two interacting cells will just touch; when closer than the equilibrium distance, the particles repel, and when further apart, they attract. The spring constant determines the relationship between the strength of the force and the distance between the particles, the magnitude of the force increasing linearly with the distance from equilibrium; thus the spring constant determines in the case of a repulsive force, the cells’ compressibility; and in the case of an attractive force, the combined effect of the strength of adhesion and the cells’ resistance to stretch. The spring constant is an adjustable model parameter, which we have tuned to optimize the model behavior, as described under Metrics: Experimental parameters, below.

Tissue Forge offers two ways to define how potentials are applied to particles. First, they can be applied to specific particle pairs, referred to as a bond. We use this approach to model cell adhesion. Two cells at a distance, which are not bonded to one another, will not attract one another. But once a pair of nearby cells becomes linked by an explicit bond, any force tending to separate those cells will be opposed by the potential’s attractive force, representing the mechanical coupling of adhesive junctions. If the balance of all these forces results in pulling the cell centers further apart, then the attractive force increases with distance, as for a spring under tension, thus limiting the separation of those cells and maintaining tissue cohesion. Alternatively, using a global approach, Tissue Forge supports applying a potential to two particle types, such that it will act the same way on any pair of particles that are members of those types, whether or not an explicit bond is present. We use this global approach to define repulsion between EVL cells, so that volume exclusion operates on nearby cells regardless of whether they are adhering (explicitly bonded). We also use this approach to define a resting distance between all EVL cells and the yolk cell center to provide basic containment to the yolk cell surface. For interactions between EVL cells, the resting distance for both attractive and repulsive potentials is based on the cell (not particle) radii as described above. In contrast, for the interaction between any EVL cell and the yolk cell, the resting distance is based on the particle (not cell) radii, so that the EVL particle surfaces just touch the yolk particle surface.

#### Forces

In addition to calculating forces based on the potentials set up by the user, Tissue Forge allows the user to apply ad hoc forces to any particle by providing a vector representation of the force, independent of any explicitly defined potential. We apply force vectors to the individual EVL margin particles to represent the driving force for epiboly, which in live embryos is generated within the yolk cell and acts only on the EVL margin. We never apply such forces to internal EVL particles; their movements are thus entirely driven by their mechanical coupling to other particles in their neighborhood.

### Initialization

We initialize our simulation by setting up a layer of EVL particles on the surface of our single yolk particle, in a configuration resembling a 30% epiboly stage zebrafish embryo. We first set the number of desired particles to 660, the reported average number of EVL cells in live embryos at 30% epiboly [14] (Table 3). From that, we calculate the cell radius to fit those particles into the available embryo surface area at that stage. First, we generate a larger number of EVL particles, enough to cover the entire surface of the yolk with cells of the calculated size, and distribute them randomly on the yolk surface. Because this procedure does not take the bulk size of cells into account, it allows the random points to be placed arbitrarily close, generating a distribution containing random clusters of unrealistically high density. We then add, as described earlier, a repulsive potential between EVL particles so that they can push each other apart to an appropriate distance, and a spring-like potential between the yolk particle and the EVL particles, to hold them to the yolk cell surface as they spread. We allow this to equilibrate by letting Tissue Forge integrate the forces as described below, resulting in a fairly even, but stochastically arranged, distribution of particles on the surface (Fig. 2a).

Next, we add spring-like bonds between neighboring EVL particles throughout the spherical layer, with a resting distance determined by the cell size as described earlier. Each particle is guaranteed to be bound to a minimum of five other nearby particles. Bonds are allowed to cross. Henceforth, “neighbors” refers simply to particles that are bonded, regardless of distance. After an additional equilibration step, this results in a mechanically coupled network of particles, representing an epithelium (Fig. 2a).

Finally, we consider the network of particles to consist of two subsets (those above the desired EVL edge position, and those below), and use the graph theory definition of a subset boundary (the set of all particles above the dividing line, that are linked to at least one particle below the dividing line) to determine which particles will be designated as edge particles and assigned the EVL margin particle type. All the particles below the dividing line, along with their linking bonds, are then deleted, and the bonds along the edge are cleaned up to ensure a ring of margin cells, each bound to exactly two other margin cells, one on each side, leaving our final set of particles and bonds representing the EVL. We take care to maintain that topology throughout the simulation as bonds are modified. Because the positions of the particles were initialized stochastically, the number of particles in each of the two subsets is also stochastic, so the final number of cells will vary from run to run, in a stochastic distribution around the initial target of 660.

### Dynamics and events

#### Physics integration

When set in motion, Tissue Forge iterates over timesteps indefinitely until stopped, integrating all forces and updating the positions and velocities of all particles. During initialization, we use this process to equilibrate our particle positions as described above; and after initialization, it drives the rest of the simulation. (Except where noted, we stop the simulation when the mean polar angle of all EVL margin cells reaches 95% of the arc from animal to vegetal, i.e. mean *ϕ* = 0.95*π*, or 171°; equivalent to ∼99% epiboly).

Tissue Forge supports both a deterministic mode of physics update, and a stochastic mode, in which particles can be subjected to a varying random force each timestep; we use Tissue Forge deterministically, and all stochasticity is introduced via our python code. (See “Sources of stochasticity in the simulation”, below, for further details.)

Tissue Forge users can provide user-defined events, in which any additional desired manipulation of the components can be done, executed after physics integration on any desired schedule. We define an event to be executed once per timestep, in which we implement three sets of update rules: 1) updating the application of exogenously imposed force; 2) bond remodeling (the stochastic breaking of existing bonds and the making of new ones); and 3) in some experiments, cell division. Once Tissue Forge completes its integration of the physics for a given timestep, it triggers our event, which executes each of these procedures as described below.

#### Calculating exogenous force

To model the force driving epiboly, which is generated outside of the EVL cells (in the yolk cortex) and acts on the EVL margin, we apply force vectors directly to the EVL margin particles. These forces are dynamically recalculated each timestep according to changing conditions. We assume that the force pulls the EVL cells in a vegetalward direction (i.e. the force vector applied to each cell is tangent to the embryo surface at the particle’s position, and points in the direction of increasing polar angle *ϕ*). In our initial versions of the model (Fig. 5a, Movie S2), we further assume that the force per unit length of EVL edge is uniform everywhere, and is constant over time. These assumptions are consistent with a force that is generated locally and which drives the EVL to stretch toward the vegetal pole; and are agnostic regarding the mechanism by which that force is generated.

Tissue Forge forces act on particles, which are discrete and, in our simulation, non-uniformly distributed. Therefore to model a force acting uniformly along the continuous EVL margin edge, we assume that the magnitude of the force vector experienced by each cell (Fig. 2d) will be proportional to the length of EVL edge the cell occupies, L_i_. We approximate that length (as an angular distance, in radians of azimuthal arc) by assuming that the cell boundary between any pair of bonded cells lies at the angular midline between the two particles. Then the magnitude of force F_i_ to apply to cell i is just:

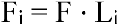

where F is a configurable parameter of the model, the force per unit length of EVL edge (Table 3, “_force_per_unit_length”). The resulting customized force vector to be applied to each particle is then passed to Tissue Forge, which adds it to all other existing forces acting on that particle during its next force integration step. In effect, the force magnitudes have been weighted according to the length of each cell’s exposed edge.

In subsequent experiments (Fig. 5b, Movie S5), we expand on this concept of weighted forces to model local regulation of the force acting on the EVL margin. Because of our approximation of the length of cell exposure to the edge, as radians of azimuthal arc, cells of a given absolute length will receive slightly more force, the further they are from the equator. This effect will be most noticeable near the pole, and may explain why the simpler approximation leads to lopsided epiboly and a small protrusion toward the pole (Fig. S1, Fig. S2, Fig. 5a, Fig. 7). To reduce this error, we now take into account the polar angle *ϕ* of each cell’s position on the embryo surface (i.e., the varying degrees of advancement of different cells toward the vegetal pole). We calculate a relative weight for each cell:

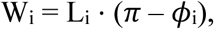

so that regions of the EVL edge that are lagging behind will be pulled more strongly than regions that are leading. We then normalize all these weights to sum to the leading edge circumference (calculated each timestep based on the mean polar angle of all EVL margin cells), and treat them as effective cell lengths, multiplying by F as above to obtain the customized force per cell.

#### Bond remodeling

On each timestep, every EVL cell then has a chance to participate in a bond-remodeling transformation (Fig. 4). In broad outline, the procedure is as follows. There are four possible transformations: any EVL cell (internal or margin) can break an existing bond to a neighbor, or make a new bond to a nearby cell to which it is not already bonded. In addition, subject to certain topological constraints, any EVL margin cell can move out of the margin into the interior of the EVL, and any internal EVL cell bound to two adjacent EVL margin cells can move between them into the margin; in either case the Tissue Forge particle type is changed accordingly. These two transformations, out of and into the margin, don’t involve actual movement, but only a change of particle type and a change in connectivity; these changes then result in movement when Tissue Forge subsequently integrates the modified forces. These margin-related transformations are handled separately from the internal ones because of special topological details at the margin, and are the only time that bonds between two EVL margin cells are created or destroyed. Each EVL cell will attempt one of these four transformations, selected with equal probability. (See “Sources of stochasticity in the simulation” for further details.) This selection can result in candidate transformations that are topologically invalid, which are disallowed; in that case no transformation takes place on that cell during that timestep. Transformations that meet the validity requirements are then evaluated according to an energetic favorability criterion, based on a preference for 60° bond angles between a cell and two adjacent bonded neighbors (Fig. 14). Breaking existing bonds, or making new ones, changes the bond angles, leading to a restoring “force” that keeps the system near that 60° configuration, by tending to reject transformations leading too far from it, and to accept transformations bringing the configuration back.

**Fig. 14.**
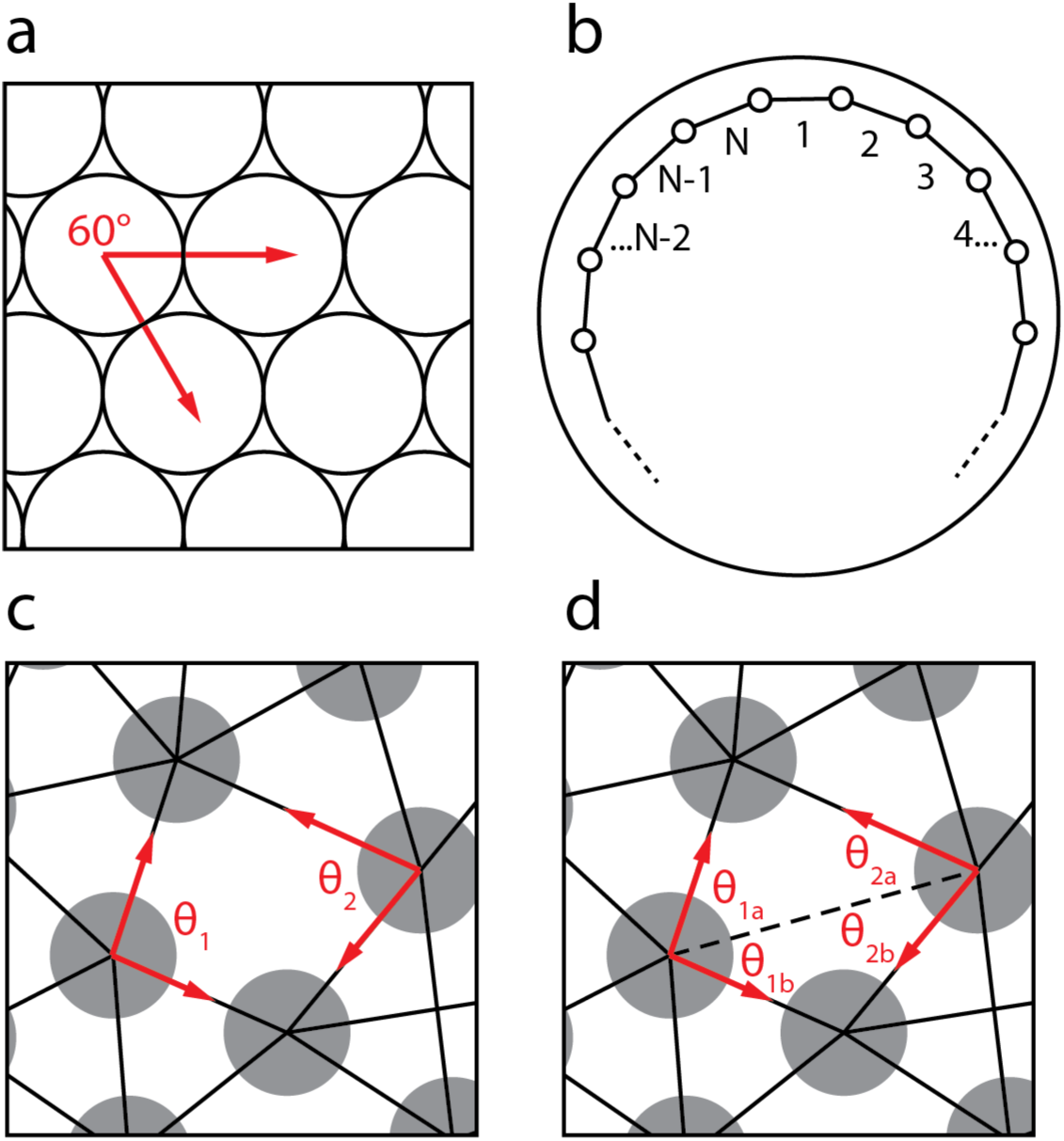
Calculating the bond angle constraint. The energy change ΔE when a bond is made or broken is a function of the actual angles between existing bonds, and a target angle. a. Everywhere except along the leading edge of the EVL, the target angle is 60° based on the angle between uniformly sized circles in a perfectly hexagonal packing arrangement. b. Schematic of the EVL leading edge as seen in vegetal view. The target angle along the leading edge takes into account the curvature of the embryo surface, and is adjusted dynamically as the leading edge contracts. The leading edge forms an irregular polygon encircling the embryo, with particles located at its vertices. Thus the number of vertices and edges (N) equals the number of particles and bonds in the leading edge; in the diagram, polygon edges are numbered. The target angle between consecutive bonds is the vertex angle of a planar, regular N-gon, equal to 180° – (360°/N). This value depends only on the number of edge particles present, and therefore decreases gradually as particles leave the leading edge during epiboly. c. In the absence of a bond between the two indicated particles, the energy depends on bond angles *θ*_1_ and *θ*_2_. d. In the presence of a bond (dashed line), the energy depends on bond angles *θ*_1a_, *θ*_1b_, *θ*_2a_, and *θ*_2b_, such that *θ*_1_ = *θ*_1a_ + *θ*_1b_ and *θ*_2_ = *θ*_2a_ + *θ*_2b_. ΔE is the difference between the energies of the two configurations (c and d); hence ΔE_bond formation_ = –ΔE_bond breaking_, and one configuration (presence or absence of the bond) will be favored over the other.

In its complete detail, the bond remodeling process at each timestep proceeds as follows:

1. Each EVL cell is visited, in random order.
2. For each visited cell, one type of transformation is selected at random for the cell to attempt. Internal EVL cells may attempt either to form a new bond with a neighbor, or break an existing bond. Marginal cells have four possibilities: form a new bond (to an internal EVL cell), break an existing bond (with an internal EVL cell), become an internal EVL cell (a new bond is formed between its two marginal neighbors), or break an existing bond to one of its marginal neighbors (a nearby internal EVL cell bound to both marginal neighbors becomes marginal). None of these transformations involve altering the position, or any other property, of any particles; they only modify the force relationships between particles, and those changes influence subsequent motion determined by the Tissue Forge force integration. The latter two transformations always involve the force relationships of three particles near the margin: one that either gains or loses margin identity, and the two margin particles on either side of it.
3. For each of these attempted transformations, neighboring partner cells for the transformation are selected; and the transformation is evaluated for validity. If found invalid, the transformation is rejected and no transformation occurs on that particle during that timestep (though the particle may be selected as a partner when one of its neighbors is visited). Specifically:

3a The visited cell attempts to form a new bond (Fig. 4a). There are no invalid scenarios in this case. The nearest neighbor cell to which the cell is not already bonded, is selected as a partner. (If the cell is a margin cell, only internal EVL cells will be selected as a partner; forming new bonds between pairs of margin cells is considered under 3c.) A bond will be created between the two cells, provided the transformation is accepted (according to criteria described below, under step 5).
3b The visited cell attempts to break an existing bond (Fig. 4a). A candidate bond to be broken is selected from among the set of all of the cell’s bonds, excluding “edge bonds” (bonds between a pair of EVL margin cells). In other words, for an internal EVL cell, all of its bonds are valid candidates to be broken; for an EVL margin cell, only its bonds to internal cells are valid candidates. (Bond-breaking between pairs of margin cells is considered under 3d.) If no valid candidates are present (the cell is a margin cell and is bonded only to other margin cells and not to any internal cells), then no transformation takes place. Otherwise one of the candidate bonds is selected at random and will be removed, provided the transformation is accepted.
3c A visited EVL margin cell attempts to become internal (Fig. 4b). The transformation is only valid if the leading edge is unambiguously concave (the position of the cell is closer to the animal pole than are either of the two neighboring margin cells to which it is bonded, hence the cell is already positioned appropriately for migration inward); otherwise no transformation occurs. If valid, and provided the transformation is accepted, the particle type of the cell is converted to internal EVL, and a bond is formed between the two neighboring margin cells. The bonds of the converting cell itself are not altered, so it remains bonded to the two marginal neighbors.
3d Upon visiting an EVL margin cell, a nearby internal EVL cell attempts to become marginal (Fig. 4b). The visited margin cell, and one of its bonded marginal neighbors (to the left or right, selected at random), define an attempted insertion site. An internal cell already bonded to both of these marginal cells is selected. If no such suitably bonded internal cell exists, then the insertion site on the other side of the visited margin cell is tried; if no suitable internal cell is found on either side, then the transformation is invalid and does not occur. If more than one such suitable cell are found on the same side, then the one closest to the two margin cells is selected. (I.e., the sum of each cell’s distance to the two margin cells is calculated, and the one with the lowest sum is selected.) The selected internal cell must then pass two additional validity checks. First, the cell must be within 0.75 cell diameters of the margin (defining the margin as the position of whichever of the two margin cells is closest to the vegetal pole, and one cell diameter as the mean diameter of all three cells). Otherwise the transformation is invalid and does not occur. Second, the selected internal cell must not be bonded to any additional margin cells besides the two at the insertion site; this restriction simplifies the computation by avoiding the topological complexity of such an edge case. If the internal EVL cell passes both of these validity tests, and provided the transformation is accepted, then the bond between the two neighboring margin cells is removed, and the internal EVL particle type is converted to marginal EVL. The bonds of the converting cell to the margin cells at the insertion site are not altered. Finally, internal EVL cells are only allowed to have bonds to at most 3 margin cells. If, by virtue of the internal EVL cell changing its type from internal to marginal, a nearby cell now exceeds that limit, the bond to that cell is broken.
4. Additional validity tests apply to all four transformation types. First, each cell is required to maintain a minimum of 3 bonds to neighboring cells. If a bond-breaking event would reduce a cell’s connectivity below that threshold, then the event is rejected. Second, if a bond-making event would result in an internal EVL cell being bonded to more than 3 margin EVL cells, then the event is rejected.
5. Finally, transformations that pass their validity checks, regardless of transformation type, are accepted or rejected according to an energy minimization criterion in the form of an elastic constraint. The energy of a given system configuration (the positions of all cells and bonds) is a measure of its departure from a target configuration, just as the energy of a bond is a measure of its departure from its resting length. The target configuration represents the lowest possible energy state and is defined in terms of a preferred bond angle between consecutive bonds on the visited cell, as follows. For all bond angles involving at least one internal EVL cell, the target angle is 60° (Fig. 14a). For the angle between three consecutive margin cells (i.e. the angle along the leading edge of the EVL, ideally “straight” to the extent allowable on the curved embryo surface), we determine its target value based on the polygon that would be formed by the current number of margin cells, all lying in a plane (Fig. 14b). If N is the current number of margin cells, then the target angle will be the average vertex angle of that polygon:

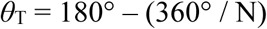 This value is thus largest (approaching 180°) when the cell population of the margin is largest, near the embryo equator; and decreases as the margin population shrinks, as the leading edge approaches the vegetal pole. When a bond is formed or broken, the energy of the system changes. If the change is favorable (negative; the transformation moves the system configuration toward its target), the transformation is always accepted. If the change is unfavorable (positive; the system configuration moves away from its target), the transformation is accepted with a probability that decreases as the energy increases. Thus the configuration is free to move to higher energy states locally, but the greater the departure from the target, the lower the probability of that occurring. Specifically, to calculate the energy change associated with the making or breaking of a bond, we first calculate the energy before and after the transformation, based on the bonds that are present, and the angles between adjacent pairs of bonds, as labeled in Fig. 14. Thus the energy when the bond is absent (Fig. 14c) is:

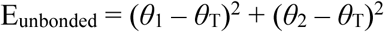

where *θ*_T_ is the target angle of 60° (except along the margin edge, where *θ*_T_ is closer to 180°, calculated as described above). When the bond is present (Fig. 14d), the energy is:

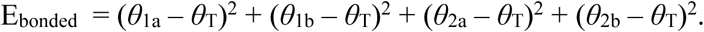 Bond breaking and bond formation are thus symmetrical, with opposite energies:

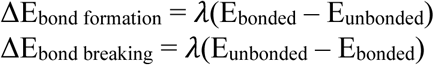

where *λ* is the strength of the constraint, a configurable parameter in the model (specified independently for the bond angles internal to the EVL and those along the edge; Table 3, “lambda_bond_angle” and “lambda_edge_bond_angle”). Then if ΔE ≤ 0, the transformation is always accepted; and if ΔE > 0, the transformation is accepted with a probability

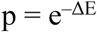 Thus transformations that are slightly energetically unfavorable (ΔE positive and small) will often be accepted, while transformations that are very unfavorable (ΔE positive and large) will be much less likely to be accepted. In this way, cell-cell connectivity can explore alternative configurations but will tend toward the energetically favorable target values, contributing to tissue stability.

#### Cell division

When the simulation is initialized, a cell division rate is calculated based on two configuration values: the developmental stage (percent epiboly) when cell division should cease, and the total number of cell divisions that should occur by that stage. (Thus the instantaneous division rate is constant with respect to EVL area increase rather than to time.) In conformance with [14], we configure cell division to cease at 55% epiboly, and to generate around 412 divisions by that time, or a 62% increase from the ∼660 cells present at 30% epiboly (Table 3). On each timestep prior to cessation, we calculate an expected number of divisions based on the division rate and the EVL area increase since the previous timestep, and an actual number of divisions from a Poisson distribution around the expected value.

Each cell in the embryo is allowed to divide at most once during the simulation, so the calculated number of dividing cells is selected at random from among all the as-yet-undivided EVL cells. The rationale for this constraint is that although cell cycles within the EVL are asynchronous, we assume that they are roughly comparable in length; the range of cell cycle lengths likely varies over less than a factor of 2, so we do not expect any cell to divide twice, before other cells have divided even once. This assumption simplifies computation because it allows us to make other assumptions about cell size: since the cell radii are uniform at initialization, there will only be two possible cell sizes once cells begin dividing: larger cells (undivided), and smaller cells (divided once). The number of cell divisions configured is not enough to exhaust all the undivided cells, so some remain undivided through the end of the simulation.

We then split each dividing cell into two daughter cells placed side-by-side. The division axis is randomly oriented within the plane of the epithelium (except for dividing margin cells, which for computational simplicity always divide horizontally, so that the two daughters are both margin cells, positioned at the same polar angle *ϕ* as the parent; they are free to migrate out of the margin subsequently if the local forces so dictate; see “Bond remodeling”, above). The daughters are each half the volume of the original, and since these are squamous cells dividing within the plane of the epithelium, their thickness (not explicitly represented) is assumed to remain constant; therefore their apical surface area is half that of the parent, so each daughter is assigned a new cell radius equal to that of the parent multiplied by 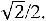 The “mass” property of each daughter (representing the drag coefficient; see “Model objects”, above) is likewise set to that of the parent multiplied by 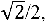 because the drag experienced by a squamous cell of uniform thickness within an epithelium should be proportional to its lateral surface area (i.e., excluding apical and basal) and hence to its circumference, so when cell volume changes during division, drag should scale as a one-dimensional property, like radius.

This division operation is carried out by the Tissue Forge “split” function, which leaves the parent particle’s bonds intact, and creates a second identical daughter particle with no bonds. We set the new cell radius for each particle as described above. (Tissue Forge also automatically halves the particle volumes. We override that and keep particle radius constant; only cell radius changes during division.) We then update the existing bonds on the parent particle to reflect their dependence on the now modified cell radius, and we add bonds connecting the new daughter with its nearby neighbors, using the same procedure as during initialization of the simulation (though in this case requiring only a minimum of three bonded neighbors rather than five).

### Sources of stochasticity in the simulation

The following events during the execution of the simulation are determined stochastically:

Initialization:

- Determining exactly how many particles of each type to create, and their positions, as described under Initialization, above.

Bond remodeling:

- Each timestep, each cell is visited and has an opportunity to attempt a bond transformation, but the cells are visited in random order, and the order is different each timestep.
- Selecting which transformation (making/breaking bonds; entering/leaving the margin) is attempted by each cell: a transformation type is selected with equal probability from the possible transformations on each EVL cell, depending on its type (internal or margin).
- When we attempt to break one of a cell’s bonds, the bond to be broken is selected at random from the valid candidate bonds on that cell.
- When we visit a margin cell and attempt to find a nearby internal EVL cell able to enter the margin next to it, we make that attempt to the left or the right of that margin cell with equal probability. (If a suitable internal cell isn’t found on that side, we try the other side.)
- Energetically unfavorable transformations (energy of the transformation ΔE > 0) are accepted with a probability equal to e^-ΔE^, that is, with vanishing probability as the energy increases.

Cell division:

- The number of cells that divide in a given timestep is determined by a Poisson distribution around a value reflecting the amount of area increase of the EVL since the previous timestep. In other words, the division rate is calibrated to the area increase, so the overall rate is determined by two configuration parameters: at what stage cell division should cease, and how much the EVL cell population should increase by that time (Table 3, “cell_division_cessation_percentage”, and “total_epiboly_divisions”). The target rate per unit of area increase is thus constant; but at each timestep the number of cells that actually divide varies stochastically around that target.
- Once the number of cells to divide in a given timestep has been determined, those cells are selected at random, from among all the EVL cells that have not yet divided.
- Interior EVL cells divide with their division axis oriented randomly within the sheet. (For computational simplicity, EVL margin cells always divide horizontally.)

### Metrics

#### Basic units

Units of distance, time, force, and mass (or drag coefficient; see “Model objects”, above) are arbitrary in Tissue Forge. We have used a diameter of 6 distance units for our yolk particle, and 0.09 distance units for the particles making up the EVL. Thus our embryo diameter is 6.18 units. A live zebrafish embryo has a diameter of 0.7 mm [4], so one Tissue Forge distance unit in our model represents about 0.113 mm. Our simulations spanning from 30% epiboly to 99% epiboly, a period of about 5 h 20 m in live embryos [4], vary in elapsed time between around 2,600 and 4,000 timesteps, so one timestep represents about 5-7 s of development time. We set the mass (drag coefficient) of EVL cells to 1.0.

#### Experimental parameters

Three aspects of the model – cell adhesion, the bond angle constraint, and the vegetal pulling force – depend on tunable numerical parameters, which we optimized for performance by trial and error (Table 3). Their optimized values depend on and thus reflect the arbitrary basic units of distance, time, and force. At extremely high or extremely low values of these parameters, the simulation fails, but epiboly is robust to significant variation in these values, allowing for interesting experiments (Figs. 10, 11).

Cell adhesion: the spring constant of a spring is a measure of the amount of force required to stretch or compress the spring by a given distance. We use springs to model cell adhesion, but, since the springs act on particles (the centers of mass of two bonded cells), the spring constants actually represent the overall mechanical properties of adhering cells as a system: encompassing not only the strength of adhesion itself, but also the stiffness of the cells. Optimal value was 0.5. (See Fig. 11 for variations.)

Bond angle constraint: the constraint strength *λ* determines the strictness of the energy minimization criterion for rejecting provisional bond remodeling events. Optimal value was around 3.75. For internal bond remodeling events (not involving bonds between two edge particles), we held that value constant. For edge remodeling events, we used 4.0, which was convenient when experimentally varying the value (see Fig. 10).

Force: Throughout, we used 0.7 force units per unit of length of leading edge. In Model 1, this force was applied uniformly. In Model 2, this value determined the total amount of force applied over the whole leading edge, but we weighted it locally according to the position (polar angle) of each edge particle. (See “Calculating exogenous force”, above.)

#### Epiboly stages

Stages of epiboly are conventionally described in terms of percent-epiboly, defined as the position of the EVL leading edge, along a straight line from the animal pole to the vegetal pole [4], which is equivalent to the fraction of the embryo surface area covered by the EVL. From the perspective of positions on the embryo surface, this measurement becomes increasingly imprecise near the vegetal pole; a positional difference of a given % epiboly represents ever greater surface distances as one approaches the pole. We use instead, both within our code and in all of our figures except where noted, the polar angle *ϕ*, which describes particle position along a longitudinal arc from animal to vegetal along the embryo surface. The position of a particle is the position of its center of mass; we measure leading edge position for the embryo as a whole, as the mean polar angle of all of the leading edge particles. This measurement captures position with the same precision whether at the vegetal pole, or at the equator. (Polar angle also has the advantage that it can be used more generally, as one of the spherical coordinates of any cell within the EVL, and to describe the changing position of any point in the EVL as the tissue expands vegetally.) However, we configure certain options in our simulation (where to place the leading edge when initializing; at what stage cell division should cease) using percent-epiboly, and convert those values to polar angle using the formula:

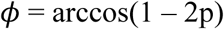

where p = percent-epiboly expressed as a value between 0.0 and 1.0. For labeling the stage of rendered screenshots in our figures, we convert back to % epiboly.

We note for the sake of accuracy that the term “30% epiboly”, referring to the first stage of epiboly after dome stage (the starting point for our simulations), should not be taken as a literal quantitative description. Initializing our simulation with p = 0.30 resulted in a configuration not resembling any published photo of a “30% epiboly” embryo, having a leading edge much too close to the animal pole. By trial and error, the value resulting in the familiar configuration turned out to be p = 0.43. It is commonly understood what a “30% epiboly” embryo looks like; however, this phrase is not a measurement, but a name, and the stage customarily referred to by that name is actually 43% epiboly, using the quantitative definition – as can be confirmed by direct measurement of the canonical photo of a “30% epiboly” embryo in [4], which described the blastoderm margin at this stage as being 30% of the way between the animal and vegetal poles, “as one estimates”. We initialize our simulation using p = 0.43 (Table 3), but throughout this paper follow the usual convention of referring to the stage that represents as 30% epiboly. We refer to all other stages according to their quantitative measurement.

#### Timecourse plots

We plot most metrics against leading edge position (polar angle *ϕ*) as a proxy for time, representing epiboly progress. In Fig. 6a, we show the relationship, as a consensus over many simulation runs, between polar angle and normalized time. Except where noted, simulations are terminated when the mean polar angle of all leading edge particle positions reaches 0.95*π*, or ∼99% epiboly; total time elapsed varies significantly among individual runs, ranging from 2600 to 4000 timesteps for our standard model (and can be further affected by alterations in physical parameters); the time axis of each run was then scaled so that its final timestep corresponds to time 1.0, before calculating the consensus plot. Simulations running Model 1 don’t reach our completion criterion for mean polar angle because the EVL projection necessitates an earlier termination; we compensate for this in plots against normalized time by adjusting the time scale to map the final timestep to the median fraction of total timesteps completed (∼0.8, Fig. 6a).

For all consensus plots of data from multiple simulation runs (Figs. 6, 8, 10, 11), we resample the datasets for each run of a given treatment to align their x-coordinates, and then plot, for each position along the horizontal axis, the median data value over all the runs in that treatment.

#### Speed of leading edge advancement

Speed of the leading edge is calculated as the change in leading edge position (see Epiboly stages, above) over time, as radians per simulation time unit.

#### Tension

Tension is a force, thus a vector quantity. For our measurements, we capture a scalar representation of the tension of an individual bond:

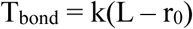

where k is the bond’s spring constant, L is the length of the bond (the distance between the centers of the two bonded particle), and r_0_ is the resting length of the bond. Thus T_bond_ is a signed scalar quantity, positive for a bond under tension and negative for a bond under compression. To obtain a single value representing the average circumferential tension of the model embryo (the tension at the EVL leading edge) at a given point in time, we simply measure T_bond_ for all the bonds along the EVL edge, i.e. those bonds connecting two edge particles, and compute their median value.

#### Margin lopsidedness

Margin lopsidedness is a measure of whether the leading edge has proceeded asynchronously (faster in some regions of the edge than in other regions). Under perfectly synchronous epiboly, lopsidedness will be near zero. To calculate lopsidedness, we first determine the best-fit plane to the positions of all the leading edge particles. Since the embryo position in simulation space is fixed, with the animal pole coordinates held directly above the vegetal pole coordinates (the animal pole has a greater z, and the same x and y, as the vegetal pole), the best-fit plane will be close to horizontal in a synchronous embryo, and tilted to some greater degree in a lopsided one. We calculate the direction of the upward-pointing normal vector to this best-fit plane, expressed in spherical coordinates, and take its polar angle *ϕ* as the measure of lopsidedness. Thus the theoretical range of the measurement is between 0 (not lopsided) and *π*/2 (if the plane were tilted all the way to vertical). In practice we see values for synchronous embryos less than around 0.03*π*, and for lopsided embryos as high as around 0.09*π*.

#### Straightness Index

To quantify the straightness of the EVL margin, we use the Straightness Index [19,20], modified for use on a spherical surface. Straightness Index (SI) for an open path through 3D space is the ratio of the “beeline” distance or shortest path between two points, to the length of the actual path traversed. (Thus SI ranges from 0 to 1, with 0 representing an infinitely tortuous path, and 1 representing a perfectly straight path.) Complications arise from the fact that we are measuring a closed path, and from the curvature of the embryo’s spherical surface to which our path (and hence the beeline path) is confined. To calculate SI in this context we first generate a cylindrical projection of the embryo surface by mapping the positions of the EVL leading edge particles to a Cartesian coordinate system whose axes are the particle spherical coordinates *ϕ* and *θ*. We then calculate the SI of the particle path within the projected flat surface, measuring distances in radians. The beeline distance is thus always 2*π*; and the actual path length traversed is the sum of the distances d between successive pairs of bonded margin particles, each segment length calculated from the angular difference in the particle positions:

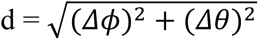

SI compares the traversed path to a reference beeline path, which in the above calculation is assumed to be horizontal. But if the margin becomes lopsided, as in Model 1, this horizontal path becomes less appropriate as the reference for comparison and would result in an underestimate of straightness. To account for this lopsidedness, we first calculate the best-fit plane to the set of margin particle positions, which will deviate from horizontal in proportion to the lopsidedness of the margin, as described above. The appropriate reference path lies within that plane. We then generate a transformation matrix to rotate the coordinate system so that the plane is horizontal, adjusting the particle coordinates accordingly, and can then calculate the SI as described above.

Together, these definitions of lopsidedness and Straightness Index allow us to measure the straightness and the lopsidedness independently; that is, to reliably assess straightness regardless of the lopsidedness of the margin, and vice versa.

## Acknowledgements

This work is based upon efforts supported by EMBRIO Institute, contract #2120200, a National Science Foundation (NSF) Biology Integration Institute. JAG and TJS acknowledge funding from NIH U24 EB028887. JAG acknowledges funding from NSF 2303695. TJS acknowledges funding from NSF 2000281. SM would like to acknowledge her father, Les Minsuk, who encouraged her interest in science at an early age, and during the writing of this paper, but who did not live to see this work completed.

## Software

The code for our epiboly simulation is available from https://github.com/sminsuk/Epiboly. The specific version used to generate all results in this version of the manuscript is https://github.com/sminsuk/Epiboly/tree/bioRxiv-release-0.2.

Running the simulation requires installing the Tissue Forge simulation environment, available from https://github.com/tissue-forge/tissue-forge [18]. We used Tissue Forge version 0.2.0.

## Appendix Epibolic deformation of the zebrafish EVL is Convergent Extension

The term “convergent extension” has been used inconsistently in the literature; for the purposes of this study we define it (after [24–26]) as a tissue-level rather than a cell-level phenomenon; and as a tissue shape change rather than the cellular mechanisms underlying it; and therefore, not as a cause of morphogenesis, but as a macroscopic description of it. The two components of CE – convergence, and extension – can also be combined with deformation in the third dimension: either thinning or thickening (e.g. CE of the neurectoderm during gastrulation and neurulation in the frog *Xenopus laevis* [25]), and can be deployed independently of one another (e.g. convergent thickening, without extension, in *Xenopus* somites [25] and in zebrafish neurectoderm [27]; convergence alone in zebrafish lateral mesoderm [27]). A successful model of zebrafish epiboly will produce the observed tissue-level change and enable exploration of alternative cellular mechanisms. When a tissue changes shape, the cells within the tissue come to rest in new positions, but CE does not imply a particular cellular mechanism: it may involve cell rearrangement or cell shape change or a combination of both, and any cell movements or shape changes may be endogenously driven by the cells of the elongating tissue, or imposed on them by forces exerted on the tissue by neighboring tissues. In that context, when we speak of convergence of cells, we refer simply to their movement to arrive at the new position, not to mechanism. In any given model system, the cellular and molecular mechanism(s) of CE must be elucidated empirically, e.g. [26,28], and can vary even among different regions of the same embryo, as in the zebrafish deep layer [27]; indeed, between different regions of the same cell [29–31].

Notably, zebrafish epiboly involves two distinct convergent extension processes, of two different types. Classically, convergent extension refers to the directional convergence of cells, moving laterally toward the midline of a tissue, while that tissue extends along its midline. Zebrafish gastrulation, which takes place between 50% and 100% epiboly (tailbud stage), involves the convergence of lateral cells toward the dorsal midline, and the narrowing and elongation of dorsal axial tissue – convergence and extension of the classical type [16] (Fig. 1b in the main text). Inherently, this type of CE requires either a pair of free lateral edges that approach each other, such as in an explant [32], or differential thinning, where the ventral part of the tissue becomes depopulated in order to feed cells to the dorsal part (“dorsal convergence and ventral divergence” in *Xenopus* [25]; the “evacuation zone” in zebrafish [4,33]). Axial CE in zebrafish involves the deep cell layer and even the YSL; the EVL is the only part of the embryo that does not participate in it [33–36]. The shrinking of the leading edge of the vegetally extending blastoderm during epiboly represents a different variation on convergence and extension (and thinning [14]), which occurs when a circular or cylindrical tissue shrinks its circumference uniformly as the tissue elongates; then there is neither an identifiable midline, nor a lateral edge, and cells come together along the shrinking circumference, without lateral migration (Fig. 1c in the main text). The cells may jostle around each other but net cell movement is in the direction of tissue elongation. In this mode of CE, the continuity of the deforming tissue around a central axis results in the narrowing of a cylinder, as in the secondary invagination and elongation of sea urchin archenteron [17,37]; the closing of a circular opening, as in vestibule closure during metamorphosis of the sea urchin *Heliocidaris erythrogramma* [38]; or the shrinking leading edge circumference of all participating tissues in zebrafish epiboly (including the EVL), after they pass the equator. Our model considers only epiboly and not axial development, and only the EVL, so our discussion of convergent extension refers exclusively to this second process.

## SUPPLEMENTARY MATERIAL

### Legends to Supplementary Movies

Movie S1. The model EVL, being stretched around the yolk from its initial configuration by an external force, without adding dynamic bond remodeling; mechanical coupling relationships between cells are fixed. Because individual bonds are elastic and neighbor relationships are fixed (precluding any possibility of cell rearrangement), the global behavior of the cell layer is likewise elastic. Note also the crumpling of the edge of the sheet (yellow particles) from its relatively straight path at the start of the simulation, to a highly folded structure at the end. After fully stretching the tissue around the yolk (at 0:08 in the video), we release the external forces, and the tissue deformation reverses, demonstrating the tissue elasticity. In contrast, a model of living tissue must be able to undergo permanent viscoelastic deformation, and cell rearrangement. (Tissue Forge has a programmable camera; in this and most subsequent movies, we have it automatically begin rotating to a vegetal position when any point on the EVL margin reaches a mean polar angle *ϕ* = 0.75*π*.) (Corresponds to Fig. 3 in main text.)

Movie S2. Model 1 generates epibolic movement resembling a living zebrafish embryo. However, epiboly progress becomes asynchronous at later stages, with the leading edge of the EVL shifting off-center relative to the vegetal pole, and finally ending in a protrusion toward the pole on one side. (Corresponds to Fig. 5a in main text.)

Movie S3. Model 1. Cells that have undergone division are labeled brown if located in the EVL margin, and light blue if located in the EVL interior. (Corresponds to Fig. S1.)

Movie S4. Model 1 with cell division disabled. (Corresponds to Fig. S2.)

Movie S5. Our enhanced, regulated model (Model 2), in which externally applied forces on the EVL leading edge are adjusted locally to ensure epiboly progresses synchronously around the whole embryo. (Corresponds to Fig. 5b in main text.)

Movie S6. Lineage tracing. Model 2 with EVL margin cells labeled red at simulation initialization; all cells retain their color through the entire simulation. Cell division was disabled; therefore the observed widening of the labeled tier and the cell mixing are not due to cell division but only to cell rearrangement. (Corresponds to Fig. 9a in main text.)

Movie S7. Lineage tracing, set up as in Movie S6, but labeling tier 1 of internal EVL cells. (Corresponds to Fig. 9b in main text.)

Movie S8. Lineage tracing, set up as in Movie S6, but labeling tier 5 of internal EVL cells. (Corresponds to Fig. S3.)

Movie S9. Lineage tracing, set up as in Movie S6, but labeling a patch of EVL cells on one side of the embryo, adjacent to the leading edge. (Corresponds to Fig. 9c in main text.)

Movie S10. Lineage tracing, set up as in Movie S6, but labeling a patch of EVL cells on one side of the embryo, adjacent to the leading edge. In this instance, the patch separated completely from the leading edge. (Corresponds to Fig. 9d in main text.)

Movie S11. Model 2, with a much stronger bond angle constraint along the margin (*λ*=10). Epiboly proceeds normally but the straightening of the EVL edge is delayed until later in epiboly, and the kinks along the edge appear rigid. Compare Movie S5. (Corresponds to Fig. 10g in main text.)

Movie S12. Laser-cut experiment (removal of external stretching force), Model 2, with a remodeling-disabled phase, during which the leading edge recoils a small distance and then stabilizes for about 5s of playback, followed by a remodeling-enabled phase, during which a much larger recoil occurs.

### Legends to Supplementary Figures

Fig. S1. Epiboly progression in Model 1, with color coding added to identify daughter cells. See Movie S3. Cells that have divided are labeled brown if located in the EVL margin, and light blue if located in the EVL interior. The smaller size of the divided cells can be discerned, despite the lack of explicit cell boundaries, by their closer packing. All cell division takes place between 30% and 55% epiboly. (% epiboly stages scored and annotated, and vegetal pole marked, as in Fig. 5 in the main text.)

Fig. S2. Epiboly progression in Model 1, with cell division disabled. See Movie S4. (% epiboly stages scored and annotated, and vegetal pole marked, as in Fig. 5 in the main text.) As a consequence of stretching without cell division, individual cells acquire larger apical surface area as epiboly proceeds, as can be seen by comparing the cell packing density in the different panels.

Fig. S3. Lineage tracing, set up as in Fig. 9, but labeling tier 5 of internal EVL cells. 30% and 99% epiboly are shown. Camera rotation was disabled, to capture only the lateral view from beginning to end of the simulation, since the labeled cells never progress beyond the equatorial zone. See also Movie S8.

## References

1. Köppen M, Fernández BG, Carvalho L, Jacinto A, Heisenberg C-P. Coordinated cell-shape changes control epithelial movement in zebrafish and Drosophila. Development. 2006;133: 2671–2681. doi:10.1242/dev.02439

2. Bruce AEE, Heisenberg C-P. Mechanisms of zebrafish epiboly: A current view. Curr Top Dev Biol. 2020;136: 319–341. doi:10.1016/bs.ctdb.2019.07.001

3. Bruce AEE. Zebrafish epiboly: Spreading thin over the yolk. Dev Dyn. 2016;245: 244–258. doi:10.1002/dvdy.24353

4. Kimmel CB, Ballard WW, Kimmel SR, Ullmann B, Schilling TF. Stages of embryonic development of the zebrafish. Dev Dyn. 1995;203: 253–310. doi:10.1002/aja.1002030302

5. Morita H, Grigolon S, Bock M, Krens SFG, Salbreux G, Heisenberg C-P. The Physical Basis of Coordinated Tissue Spreading in Zebrafish Gastrulation. Dev Cell. 2017;40: 354–366.e4. doi:10.1016/j.devcel.2017.01.010

6. Cheng JC, Miller AL, Webb SE. Organization and function of microfilaments during late epiboly in zebrafish embryos. Dev Dyn. 2004;231: 313–323. doi:10.1002/dvdy.20144

7. Marsal M, Hernández-Vega A, Pouille P-A, Martin-Blanco E. Rab5ab-Mediated Yolk Cell Membrane Endocytosis Is Essential for Zebrafish Epiboly and Mechanical Equilibrium During Gastrulation. Front Cell Dev Biol. 2021;9. doi:10.3389/fcell.2021.697097

8. Cheng JC, Miller AL, Webb SE. Actin-mediated endocytosis in the E-YSL helps drive epiboly in zebrafish. Zygote. 2023;31: 517–526. doi:10.1017/S0967199423000357

9. Betchaku T, Trinkaus JP. Contact relations, surface activity, and cortical microfilaments of marginal cells of the enveloping layer and of the yolk syncytial and yolk cytoplasmic layers of Fundulus before and during epiboly. Journal of Experimental Zoology. 1978;206: 381–426. doi:10.1002/jez.1402060310

10. Betchaku T, Trinkaus JP. Programmed Endocytosis During Epiboly of Fundulus heteroclitus1. American Zoologist. 1986;26: 193–199. doi:10.1093/icb/26.1.193

11. Hernández-Vega A, Marsal M, Pouille P-A, Tosi S, Colombelli J, Luque T, et al. Polarized cortical tension drives zebrafish epiboly movements. EMBO J. 2017;36: 25–41. doi:10.15252/embj.201694264

12. Behrndt M, Salbreux G, Campinho P, Hauschild R, Oswald F, Roensch J, et al. Forces driving epithelial spreading in zebrafish gastrulation. Science. 2012;338: 257–260. doi:10.1126/science.1224143

13. Solnica-Krezel L, Driever W. Microtubule arrays of the zebrafish yolk cell: organization and function during epiboly. Development. 1994;120: 2443–2455. doi:10.1242/dev.120.9.2443

14. Campinho P, Behrndt M, Ranft J, Risler T, Minc N, Heisenberg C-P. Tension-oriented cell divisions limit anisotropic tissue tension in epithelial spreading during zebrafish epiboly. Nat Cell Biol. 2013;15: 1405–14. doi:10.1038/ncb2869

15. Keller RE, Trinkaus JP. Rearrangement of enveloping layer cells without disruption of the epithelial permeability barrier as a factor in Fundulus epiboly. Dev Biol. 1987;120: 12–24. doi:10.1016/0012-1606(87)90099-6

16. Warga RM, Kimmel CB. Cell movements during epiboly and gastrulation in zebrafish. Development. 1990;108: 569–580. doi:10.1242/dev.108.4.569

17. Martik ML, McClay DR. New insights from a high-resolution look at gastrulation in the sea urchin, Lytechinus variegatus. Mech Dev. 2017;148: 3–10. doi:10.1016/j.mod.2017.06.005

18. Sego TJ, Sluka JP, Sauro HM, Glazier JA. Tissue Forge: Interactive biological and biophysics simulation environment. PLOS Computational Biology. 2023;19: e1010768. doi:10.1371/journal.pcbi.1010768

19. Benhamou S. How to reliably estimate the tortuosity of an animal’s path: straightness, sinuosity, or fractal dimension? J Theor Biol. 2004;229: 209–220. doi:10.1016/j.jtbi.2004.03.016

20. Batschelet E. Circular statistics in biology. London: Academic Press; 1981.

21. Lemke SB, Nelson CM. Dynamic changes in epithelial cell packing during tissue morphogenesis. Current Biology. 2021;31: R1098–R1110. doi:10.1016/j.cub.2021.07.078

22. Jain A, Ulman V, Mukherjee A, Prakash M, Cuenca MB, Pimpale LG, et al. Regionalized tissue fluidization is required for epithelial gap closure during insect gastrulation. Nat Commun. 2020;11: 5604. doi:10.1038/s41467-020-19356-x

23. Holloway BA, Canny SG de la T, Ye Y, Slusarski DC, Freisinger CM, Dosch R, et al. A Novel Role for MAPKAPK2 in Morphogenesis during Zebrafish Development. PLOS Genetics. 2009;5: e1000413. doi:10.1371/journal.pgen.1000413

24. Vogt W. Gestaltungsanalyse am Amphibienkeim mit Örtlicher Vitalfärbung : II. Teil. Gastrulation und Mesodermbildung bei Urodelen und Anuren. Wilhelm Roux Arch Entwickl Mech Org. 1929;120: 384–706. doi:10.1007/BF02109667

25. Keller RE. Vital dye mapping of the gastrula and neurula of Xenopus laevis. II. Prospective areas and morphogenetic movements of the deep layer. Dev Biol. 1976;51: 118–137. doi:10.1016/0012-1606(76)90127-5

26. Keller RE. The Cellular Basis of Gastrulation in Xenopus laevis: Active, Postinvolution Convergence and Extension by Mediolateral Interdigitation. American Zoologist. 1984;24: 589–603.

27. Williams MLK, Solnica-Krezel L. Cellular and molecular mechanisms of convergence & extension in zebrafish. Curr Top Dev Biol. 2020;136: 377–407. doi:10.1016/bs.ctdb.2019.08.001

28. Keller RE. The cellular basis of epiboly: an SEM study of deep-cell rearrangement during gastrulation in Xenopus laevis. J Embryol Exp Morphol. 1980;60: 201–234.

29. Williams M, Yen W, Lu X, Sutherland A. Distinct Apical and Basolateral Mechanisms Drive Planar Cell Polarity-Dependent Convergent Extension of the Mouse Neural Plate. Developmental Cell. 2014;29: 34–46. doi:10.1016/j.devcel.2014.02.007

30. Sun Z, Amourda C, Shagirov M, Hara Y, Saunders TE, Toyama Y. Basolateral protrusion and apical contraction cooperatively drive Drosophila germ-band extension. Nat Cell Biol. 2017;19: 375–383. doi:10.1038/ncb3497

31. Huebner RJ, Wallingford JB. Coming to Consensus: A Unifying Model Emerges for Convergent Extension. Developmental Cell. 2018;46: 389–396. doi:10.1016/j.devcel.2018.08.003

32. Keller R, Danilchik M. Regional expression, pattern and timing of convergence and extension during gastrulation of Xenopus laevis. Development. 1988;103: 193–209. doi:10.1242/dev.103.1.193

33. Kimmel CB, Warga RM, Schilling TF. Origin and organization of the zebrafish fate map. Development. 1990;108: 581–594. doi:10.1242/dev.108.4.581

34. Kimmel CB, Warga RM. Indeterminate cell lineage of the zebrafish embryo. Dev Biol. 1987;124: 269–280. doi:10.1016/0012-1606(87)90478-7

35. Rohde LA, Heisenberg C-P. Zebrafish gastrulation: cell movements, signals, and mechanisms. Int Rev Cytol. 2007;261: 159–192. doi:10.1016/S0074-7696(07)61004-3

36. D’Amico LA, Cooper MS. Morphogenetic domains in the yolk syncytial layer of axiating zebrafish embryos. Dev Dyn. 2001;222: 611–624. doi:10.1002/dvdy.1216

37. McClay DR, Warner J, Martik M, Miranda E, Slota L. Gastrulation in the sea urchin. Curr Top Dev Biol. 2020;136: 195–218. doi:10.1016/bs.ctdb.2019.08.004

38. Minsuk SB, Raff RA. Co-option of an oral-aboral patterning mechanism to control left-right differentiation: the direct-developing sea urchin Heliocidaris erythrogramma is sinistralized, not ventralized, by NiCl2. Evol Dev. 2005;7: 289–300. doi:10.1111/j.1525-142X.2005.05035.x

