## Supplementary figures and images for "Modeling Epithelial Morphogenesis and Cell Rearrangement during Zebrafish Epiboly: Tissue Deformation, Cell-Cell Coupling, and the Mechanical Response to Stress"

### Supplemental Figure S1

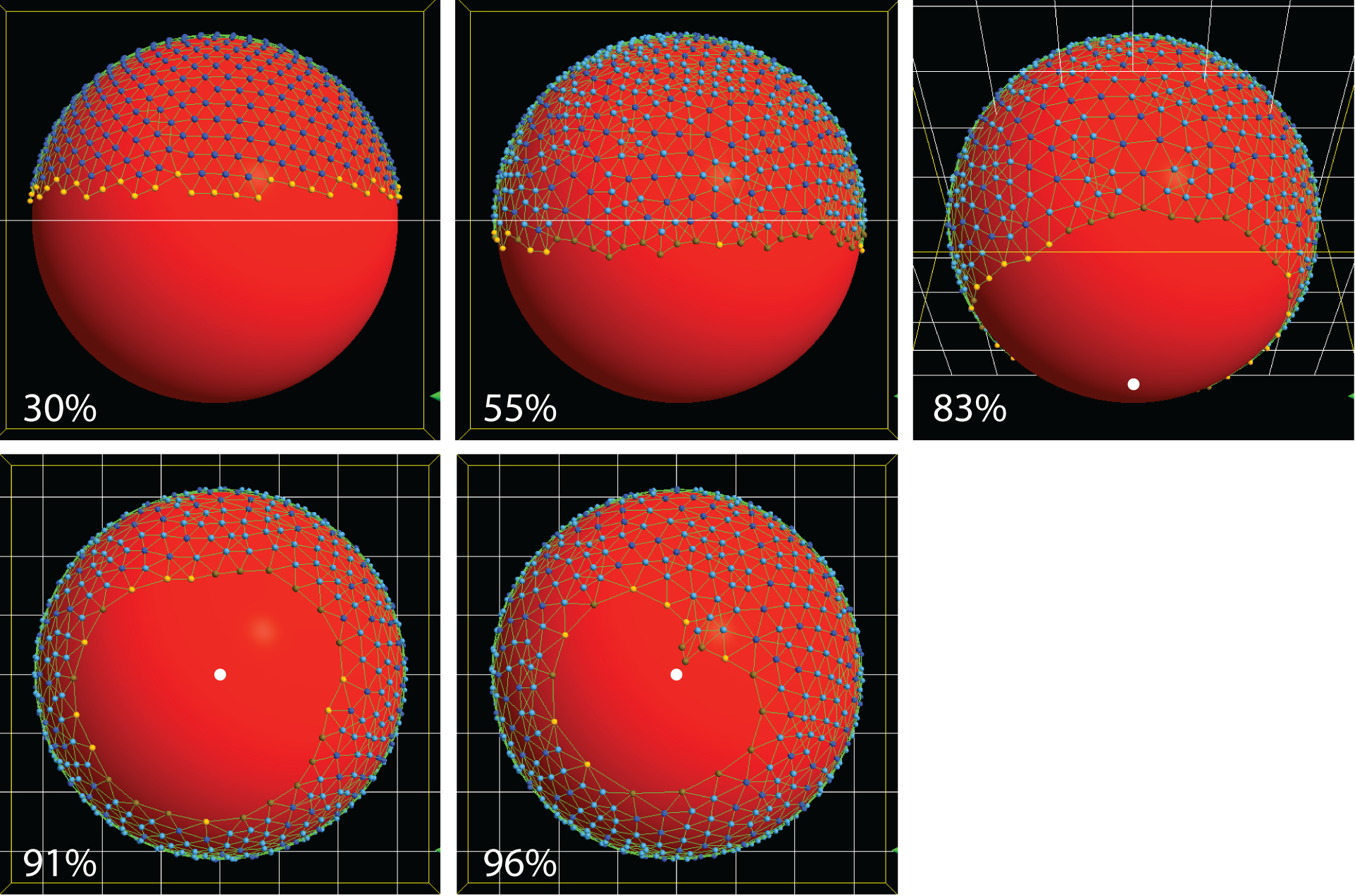

### Supplemental Figure S2

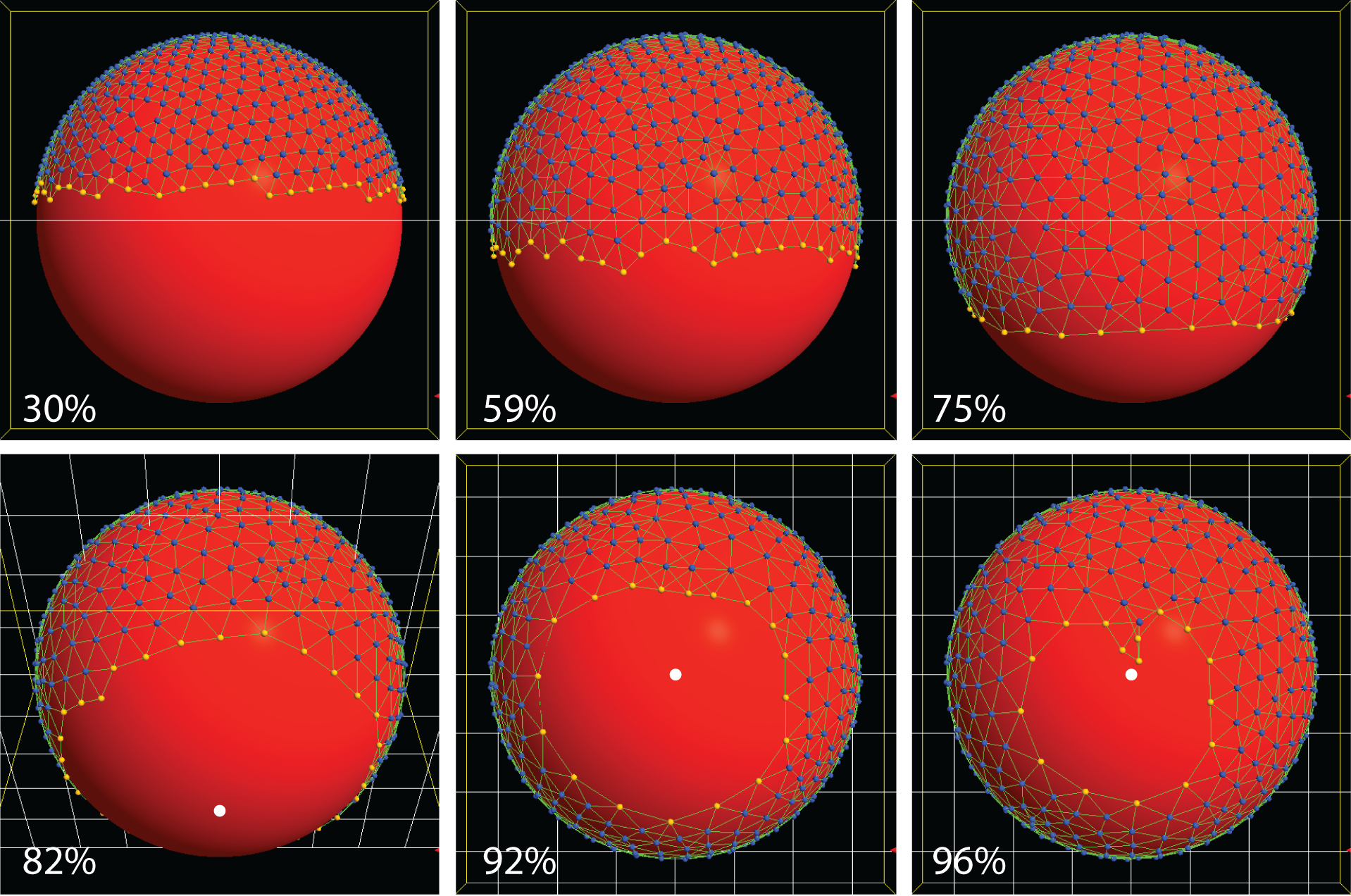

### Supplemental Figure S3

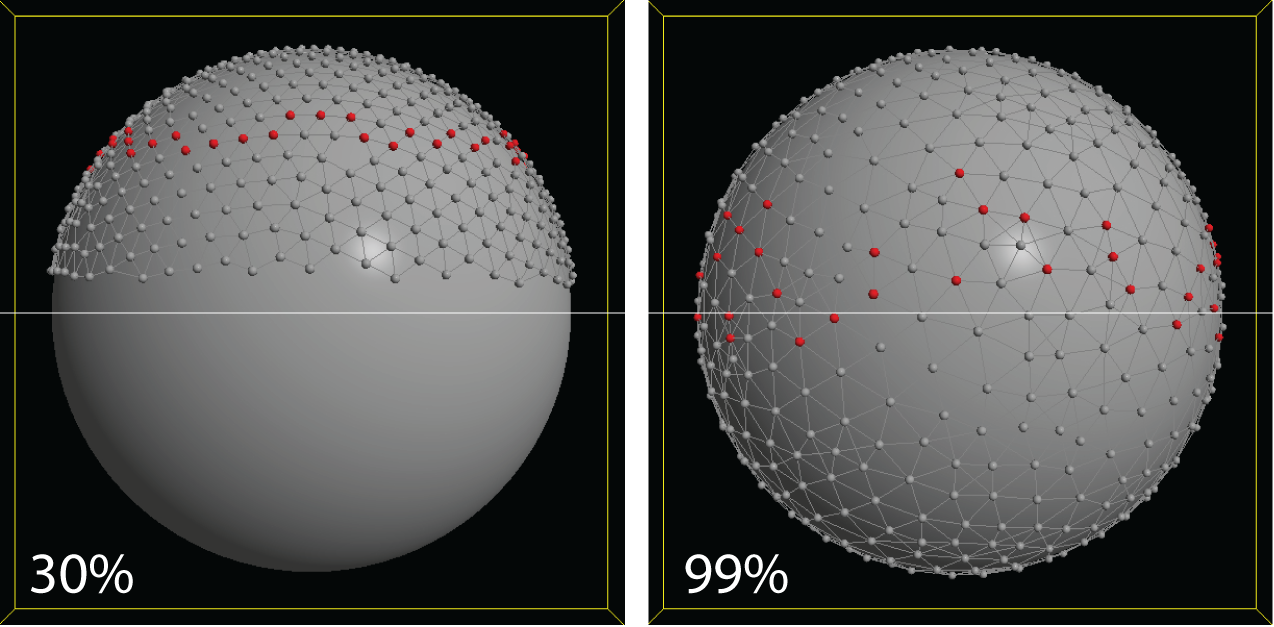
